# Bringing the Genetically Minimal Cell to Life on a Computer in 4D

**DOI:** 10.1101/2025.06.10.658899

**Authors:** Zane R. Thornburg, Andrew Maytin, Jiwoong Kwon, Troy A. Brier, Benjamin R. Gilbert, Enguang Fu, Yang-Le Gao, Jordan Quenneville, Tianyu Wu, Henry Li, Talia Long, Weria Pezeshkian, Lijie Sun, John I. Glass, Angad Mehta, Taekjip Ha, Zaida Luthey-Schulten

## Abstract

We present a whole-cell spatial and kinetic model for the ~100 minute cell cycle of the genetically minimal bacterium, JCVI-syn3A. This is the first simulation of a complete cell cycle in 4D including all genetic information processes, metabolic networks, growth, and cell division. Integrating hybrid computational methods, dynamics of the morphological transformations were achieved. Growth is driven by synthesis of lipids and membrane proteins and constrained by new fluorescence imaging data. Chromosome replication and segregation is controlled by essential SMC and topoisomerase proteins in Brownian dynamics simulations with replication rates responding to dNTP pools from metabolism. The model captures the origin to terminus ratio measured in our DNA sequencing and recovers other experimental measurements like doubling time, mRNA half-lives, protein distributions, and ribosome counts. Because of stochasticity, each replicate cell is unique. Not only do we predict average behavior for partitioning to daughter cells, we predict the heterogeneity among them.

## INTRODUCTION

To understand the rules governing cellular life, we must know how the underlying chemical and physical processes act in unison in 4D (3 spatial dimensions and time). A measurement of the cell state to quantify all chemical and physical processes simultaneously would require the exact position of and interactions between every atom inside a cell as a function of time. While such a measurement is not currently possible with experiments, strides have been made to represent complete cell states computationally.^1–4^ At the finest level of resolution, a near-atomic structure of a minimal bacterium (JCVI-syn3A) has been modeled using the coarse-grained Martini force field.^5,6^ Structural models at all-atom resolution of *Mycoplasma genitalium* and JCVI-syn3A have been created.^7,8^ Although these models achieve atomic resolutions of (nearly) complete cell states, they are either static or can only be simulated for short times (*<*1 second). Atomic-resolution models are great for capturing structure and high-resolution, short-timescale interactions, but are not capable of simulating the mechanics and chemistry that take place over minutes to hours in processes such as gene expression and cell division.

At the other end of the complexity scale, whole-cell reaction networks have been treated using steady-state flux and kinetic models. Flux balance analysis (FBA) methods have been applied to many cells from simple bacteria to eukaryotes like yeast and human cells to predict fluxes through metabolic reactions.^9,10^ FBA has proven to be a useful tool for predicting average behavior and even making claims about gene essentiality, but it lacks dynamics and cell-to-cell heterogeneity. There have been a few whole-cell kinetic models to date including *Mycoplasma genitalium* ^3^ and *Escherichia coli* ^4,11^ as well as our own model of a minimal bacterium.^1^ Although the kinetic models include reaction dynamics to dictate temporal progression of the cell state, they treat cells or cell compartments primarily as well-stirred systems lacking heterogeneous spatial organization. These well-stirred models have been proving themselves as predictive tools, but they cannot probe the dependence of stochastic processes on the spatial localization of molecular participants (e.g. RNAP must diffuse and bind to promoters on the DNA).^12–14^ Finally, generating models of cells using artificial intelligence (AI) and machine learning (ML) is a growing area of interest, and is a promising avenue for integrating the large quantities of biological data that are becoming available.^15^ However, the dynamics driving cellular life are a complicated interwoven network of chemical reactions, and there are many predictions to be made with physics- and chemistry-based models that may not be captured by AI and ML.

Building a fully physics-based model of an entire cell to explore the dynamics of cell division in a holistic manner is both highly appealing and crucial in the quest to understand the foundations of life through a bottom-up approach. In this context, an organism consisting of the fewest components and processes would provide an excellent platform. JCVI-syn3A is a genetically minimal bacterium with a synthetic genome that has been genetically reduced from *Mycoplasma mycoides subsp. capri* str. GM12.^16,17^ Syn3A has a doubling time of 105 minutes and a single 543 kbp circular chromosome consisting of 493 genes.^18^ While a previous iteration, JCVI-syn3.0, had fewer genes, Syn3A reintroduced genes allowing the cell to divide and maintain regular spherical morphologies.^17–19^ Syn3A’s minimal genome,^17,19^ abundant -omics and sequencing data,^20^ genome-wide essentiality assignment,^18^ essential metabolic map, cryoelectron tomograms,^21^ chromosome contact maps,^21^ predicted structural proteome,^22^ and now imaging of the symmetric division make it an ideal system for 4D whole-cell modeling over the entire cell cycle.

Previously we simulated the cell cycle of Syn3A as a well-stirred system using hybrid stochastic-deterministic kinetics.^1^ Several parameters in the well-stirred model were obtained from simplified 4D simulations of the first 20 minutes of the cell cycle assuming static morphology and ribosome positions from cryo-electron tomography before any DNA replication had occurred. The spatially-resolved model is more computationally expensive, but even short 4D simulations provided predictions of probabilistic factors such as the average number of active ribosomes, RNA polymerases, and degradosomes and the distribution of mRNA half-lives. From well-stirred simulations, behaviors emerged that reflected experimental measurements of Syn3A and related bacteria. For example, our model predicted the doubling time with surface area contributions arising from lipid biosynthesis and membrane protein insertion reactions.

To fully understand and characterize the spatial dynamics that dictate life for Syn3A, we need to simulate the entire cell cycle including DNA replication and dynamics, ribosome movement, and division in 4D. Here, we present the first 4D whole-cell model (4DWCM) that simulates the entire 105 minute cell cycle of Syn3A including expression of all 493 genes, kinetics of the entire metabolic network, ribosome biogenesis, chromosome dynamics including DNA replication, and morphological changes during growth and division. We again hybridize multiple simulation techniques into one model to accurately represent the broad range of lengths, concentrations, and rates that define the cell state and cell cycle progression. While the well-stirred components for metabolism remain mostly unchanged from our previous model,^1^ making the spatial components of genetic information processes and cell morphology dynamic posed significant challenges. Methods were developed to communicate a coarse-grained continuum model of the DNA to our simulation software Lattice Microbes (LM), update morphology based on the biosynthesis of membrane components, allow ribosomes to assemble and diffuse on lattice sizes smaller than their diameter, and other procedures that stem from these major additions. The DNA replication dynamics accompanied by informed kinetic parameters^23,24^from recent SMC loop extrusion experiments are in agreement with DNA sequencing data provided here. Constraints from our fluorescence imaging experiments characterizing cell morphology, DNA localization, and formation of the septum support the symmetric cell division of Syn3A. By creating a model that imposes spatial heterogeneity and its inherent stochasticity on reactions like DNA replication initiation, transcription, translation, and mRNA degradation, we uncover the dependence, sensitivity, and variations of cell cycle progression to key rates in the 4D dynamics. The 4D simulations we present here require 4 to 6 days of compute time on two high-performance computing GPUs (e.g. NVIDIA A100) per cell cycle (approximately 2 hours of biological time). Overall, we analyzed the cell cycle dynamics of 50 unique replicate cells. This 4DWCM presents a leap forward in our ability to more accurately probe the fundamental behaviors of cellular life.

## RESULTS

### Construction of the 4DWCM

Constructing a 4DWCM to simulate an entire cell cycle of Syn3A requires hybridization of computational methods at multiple levels of resolution. We have summarized the algorithm that communicates biological information between computational methods and a breakdown of the computational expenses with a flowchart shown in Figure 1. Gene expression reactions are handled with stochastic reactions and metabolism is simulated as a system of ordinary differential equations (ODEs), the details of reactions and their kinetics are all discussed in Methods. To make the chromosome(s) dynamic required the addition of yet another method: Brownian dynamics (BD) simulated on a separate GPU using LAMMPS.^6,25^ Morphological updates during growth and division are determined by the synthesis of lipids from metabolism and the translocation of membrane proteins. Each membrane component is assigned a surface area contribution, resulting in the total cell surface area growing as a function of time. For completeness, the initial counts/concentrations obtained from -omics data, a table of the kinetic parameters, and a thermodynamic analysis of the kinetic parameters are provided in Table S1 and Table S2, and the few parameter changes are discussed in Methods.

**Figure 1:**
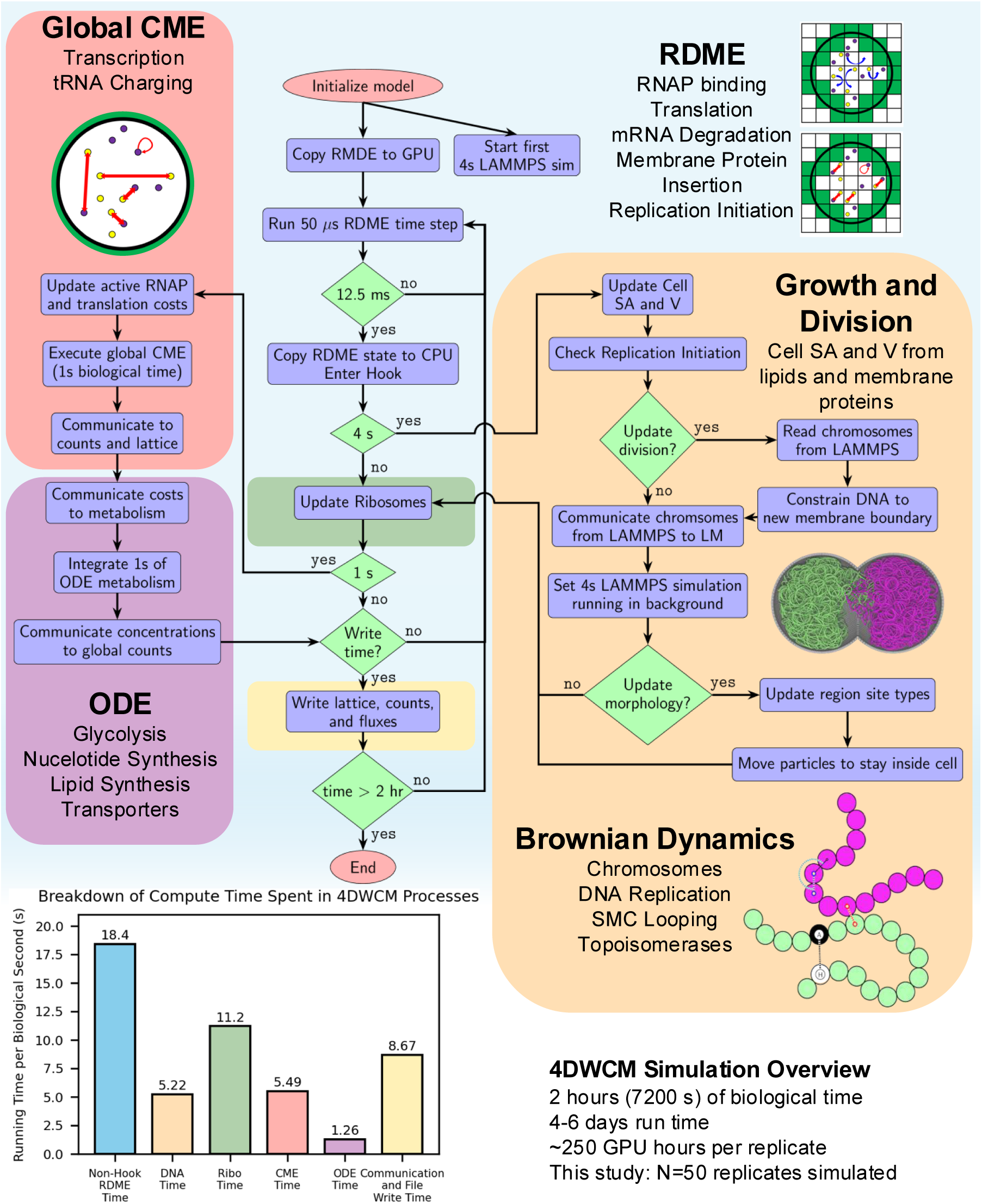
Flowchart of the hybrid simulation algorithm for the 4DWCM. The parent method is the RDME solver implemented in LM. Time steps are 50 *µ*s. The hybrid algorithm interrupts the RDME solver at an interval of 12.5 biological milliseconds. The primary methods communicated are well-stirred stochastic kinetics through a global (cell-wide) chemical master equation simulation (CME), ordinary differential equations (ODEs) for metabolism, and Brownian dynamics (integrated with LAMMPS) for the chromosomes. The average computational expense of each component are summarized in the bar chart.

### Visual representations of complete cell states

The breadth of data provided by the 4DWCM is challenging to represent visually. To summarize a few highlights of the spatial heterogeneity of our simulated cell states, we show 3D visualizations in Figure 2. Visualization of a representative trajectory following a cell through its entire cell cycle is shown in Video S1. The RDME component of the simulations requires that the 3D space be discretized to a cubic lattice, in this case with edge lengths of 10 nm. Chromosomes are represented as circular homopolymers consisting of 10 bp monomers. As lipids and membrane proteins are synthesized, we update the morphology on the lattice throughout growth and division. Figure 2A shows the progression of DNA replication, growth, and division over the course of a cell cycle. Figures 2B-G show various components of a cell state all from the same frame of time for a single cell simulation during division. All proteins and RNA are treated as individual particles as shown in Figure 2C. Chromosome dynamics are communicated to LM by imposing the LAMMPS DNA monomers onto the lattice to define excluded volume that reduces diffusion of lattice particles through and around the DNA (Figure 2D). We know genomic sequence positions down to 10 bp resolution, so we place the monomers that correspond to transcription start sites (assumed to be the first nucleotide in the respective gene’s coding sequence) onto the LM lattice. The boundary for the chromosomes in LAMMPS is smaller than the membrane boundary in LM to prevent the DNA from overlapping the membrane (Figure 2D and E). We define lattice site types with specific reaction and diffusion rules to differentiate the peripheral membrane sites from membrane and cytoplasm as shown in Figure 2F. Degradosome particles are restricted to the peripheral membrane sites, and we show the sugar transporter PtsG as a representative membrane protein that is restricted to the membrane. Ribosomes have a hybrid treatment where their centers of mass are treated as particles and their excluded volumes are treated as crosses, both on the RDME lattice as shown in Figures 2B and G, respectively (also see Figure S1).

**Figure 2:**
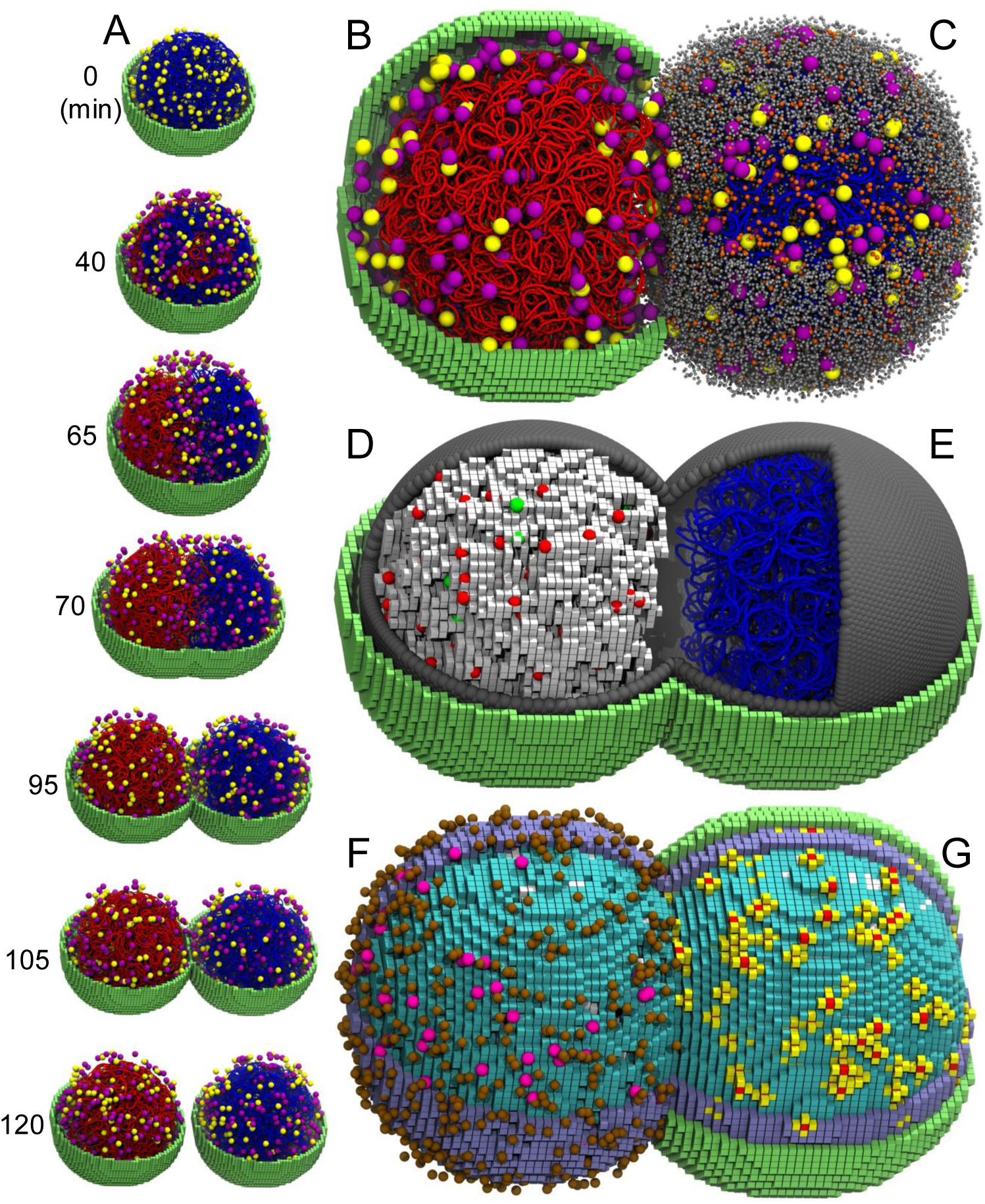
Visualization of 3D components of the 4DWCM. (A) Cell cycle progress from initial conditions to 2 hours of biological time showing lattice membrane (green 10 nm cubes), ribosomes (yellow:inactive, purple:active), and DNA (10 bp beads, blue and red differentiate daughter chromosomes). (B) Zoomed in view of the representation in A. (C) The cytoplasm is dense with proteins (grey particles) and RNA (orange particles). (D) The DNA is projected onto the lattice (white cubes) and the promoter particles (red particles:promoters, green particles:RNAP bound) are placed according to their position from LAMMPS. (E) The membrane boundary in LAMMPS (dark grey spheres) must be smaller than the lattice membrane to prevent DNA-membrane overlap on the lattice. (F) PtsG sugar transporters (brown particles) are restricted to the membrane and aren’t allowed to diffuse back into the peripheral membrane (blue cubes) or cytoplasm (cyan cubes) regions. Degradosomes (pink particles) are restricted to the peripheral membrane. (G) Excluded volume of ribosomes is maintained by projecting their volume onto lattice cubes (red cubes:center of mass, yellow cubes:excluded volume). B through G are all from the 85 minute time point from the same simulation. Generated using VMD.^53^

### Dynamics, replication, and partitioning of chromosomes

To accurately simulate the spatial heterogeneity that drives gene expression required development of a model to simulate the spatial dynamics of the chromosome configuration. This polymer model is based on our previous chromosome dynamics model.^6^ It is summarized in Figure 3 and is described in further detail in the Methods section. In our simulations we incorporate the physical effects of two classes of proteins: topoisomerases and SMC (structural maintenance of chromosome proteins, analogous to condensin). While the actions of SMC looping and topoisomerases are sufficient to substantially disentangle daughter chromosomes from one another, successful partitioning into daughter cells required the introduction of a repulsive force of approximately 12 pN force between daughter chromosomes to accelerate their movement during cell division (Figure 3A). In our simulations, DNA replication (Figure 3B) is implemented with the “train-track” model.^26^ Before DNA is replicated, the process is initiated by the essential protein DnaA. The kinetics of this fundamental process are described in detail below. To simulate the DNA’s polymer behavior, we employ an elastic worm-like chain model (Figure 3C) using the potential energy functions from Brackley et al.^27^ We also incorporate excluded volume interactions between strands of DNA (Figure 3D) and between DNA and boundary particles (Figure 3E). SMC complexes form and extrude loops of DNA, causing structural changes on the size of hundreds to thousands of base pairs per loop.^28^ As shown in Figure 3F, we mimic the effect of SMC by anchoring the loop to one bead on our homopolymer and updating the “hinge” position of the loop to progressively extrude DNA. We update anchor positions once and hinge positions ten times every 4 seconds of biological time. The rate of hinge position movement is based on experimental measurements,^24^ but it is unclear how frequently we should update anchor positions when we assume the dimers only bind to the chromosome non-specifically in Syn3A.^6,21^ Topoisomerases transiently cut DNA to allow double-strand passage and then re-ligate the DNA. We incorporate the effects of topoisomerases by periodically switching the potential from the hard excluded volume potential to a soft potential that allows DNA monomers to pass through one another as shown in Figure 3D. Without an effect to resolve strand crossings, we found previously^6^ and here that our simulations would never be able to disentangle and partition chromosomes between two daughter cells.

**Figure 3:**
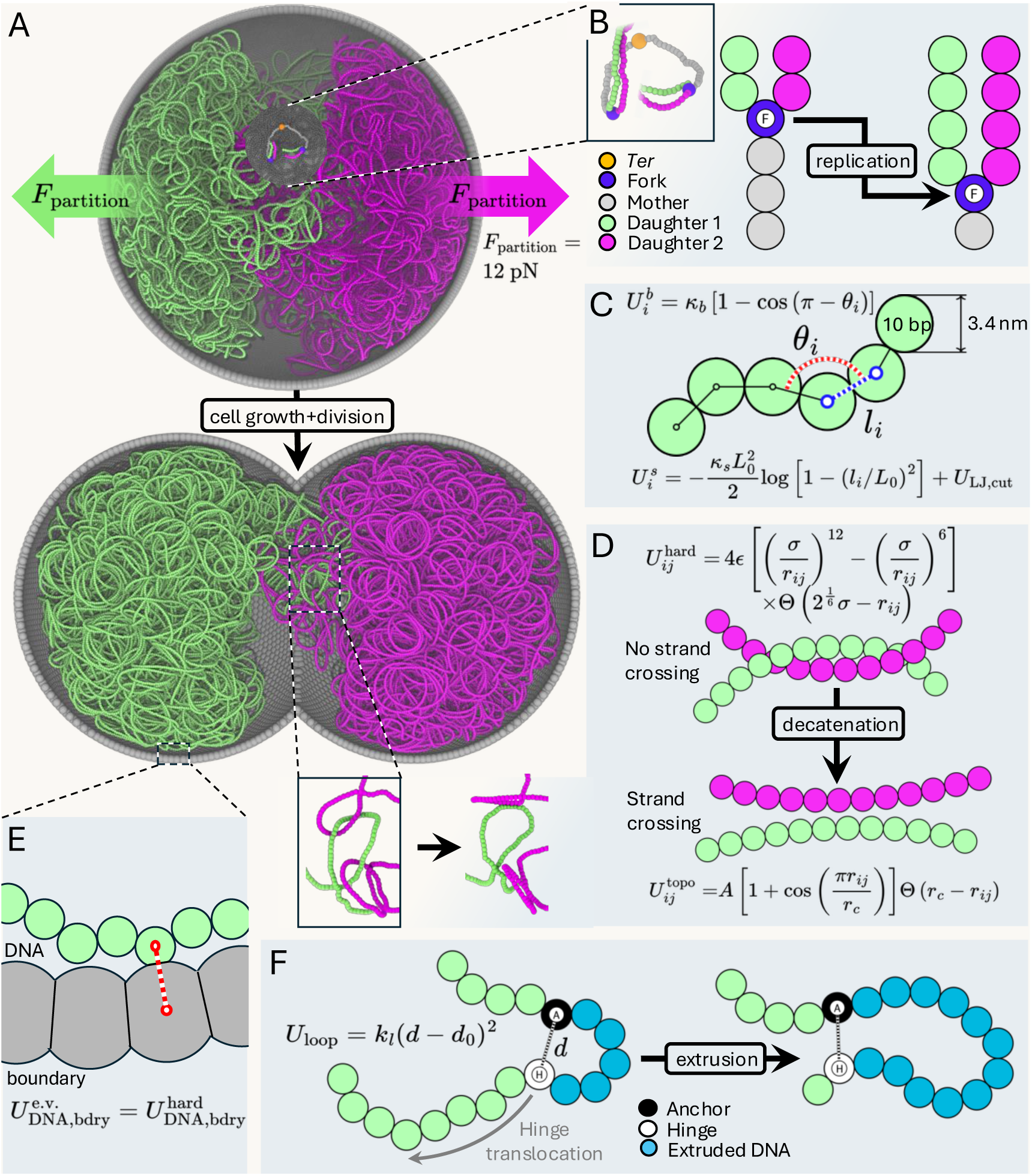
Coarse-grained modeling of the minimal cell chromosome. (A) Snapshots of the cell at approximately 45 and 90 minutes into the cell cycle. A 12 pN force is applied to each daughter chromosome in opposite directions to facilitate partitioning. A window cutout highlights the replication forks and terminus. (B) DNA replication follows the train-track model. The window cutout from panel A is magnified. (C) Bending interactions (red dashed line) impose local curvature with a persistence length 45 nm; stretching (blue dashed line) is governed by a FENE potential. DNA is coarse-grained at 10 bp resolution, with a bead diameter of 3.4 nm. (D) Excluded volume interactions between DNA strands include a “hard” repulsion to prevent strand crossing and a soft “topo” repulsion that allows it; an example where strand crossing can be seen visually is magnified. (E) Excluded volume interactions between DNA and the boundary (red dashed line) is set to be a hard-core repulsion. (F) SMC-driven loop extrusion is modeled by introducing a harmonic bond between each anchor and hinge, with hinge translocating in steps of ~20 beads simulating the reeling motion of the SMC complex. Anchors are updated (randomly placed) every 4 s; hinges are updated (translocated) every 0.4 s.

### 4D whole-cell modeling reflects experiment

In bacterial cells, the timing of the cell cycle is typically characterized by three periods for cells with a single DNA replication event: B – the time between birth and replication initiation, C – the time to replicate the DNA, and D – the time to divide the cell after the end of replication.^29^ We predict two key timings shown in Figure 4A: doubling the membrane in 105 minutes and the chromosome in 51 minutes on average. The predicted doubling time is in very close agreement with the experimental doubling time of 105 min.^18^ We made a similar prediction in our previous well-stirred model, but the predicted doubling time was 97 min.^1^ The slow-down in predicted doubling time comes from an underproduction of proteins (discussed below), resulting in slower surface area growth from incorporation of fewer membrane proteins.

**Figure 4:**
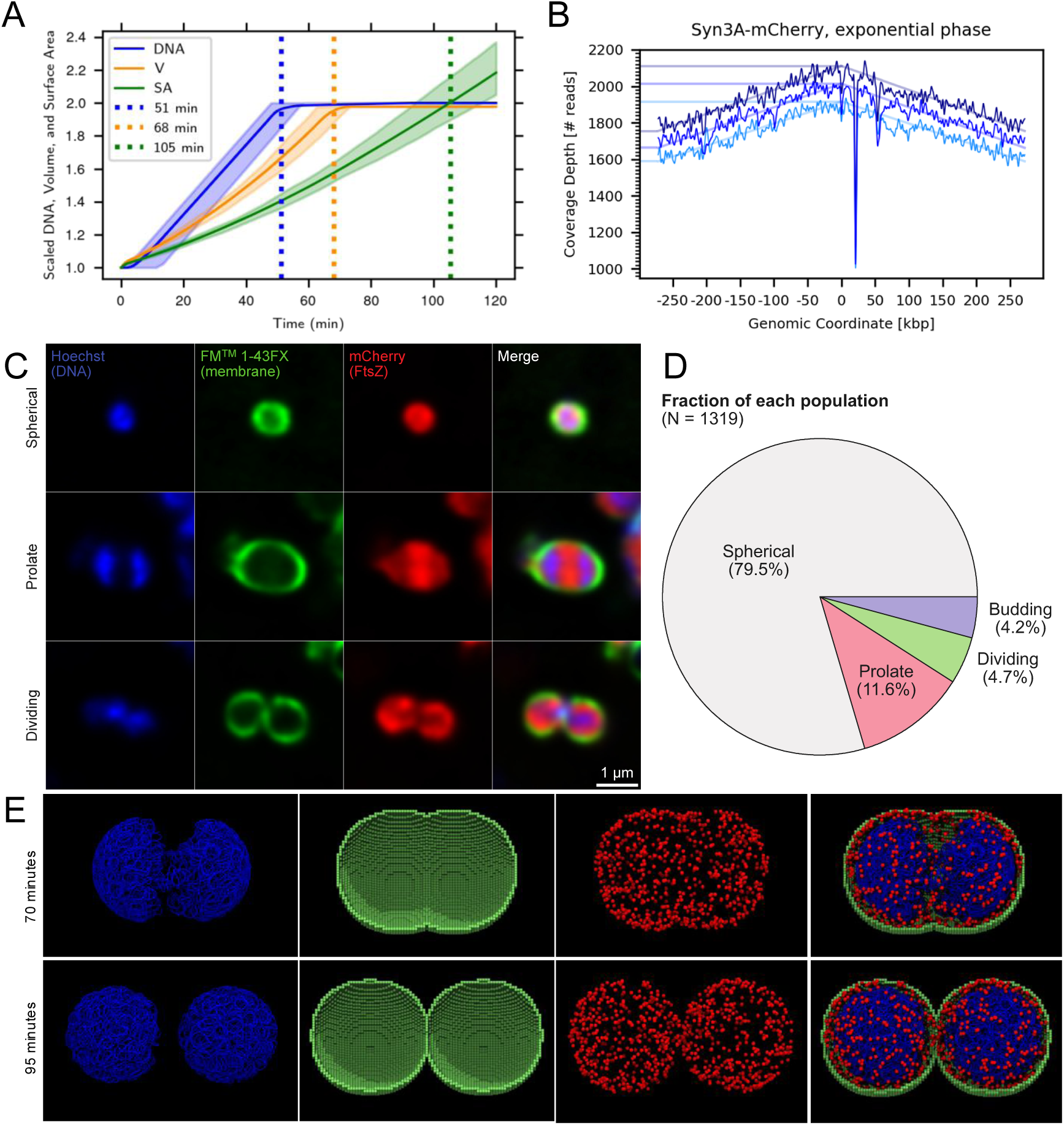
The 4DWCM accurately mimics experiments. (A) Timing of DNA, volume, and surface area doubling in simulated cell cycles. Vertical lines represent average times among the population. The combined DNA and surface area doubling times predict a ori:ter ratio of 1.28. Shaded regions represent the full range of the simulated population except for the DNA. The DNA shaded region excludes the cell with a replication initiation time of 46 minutes. (B) Experimental DNA sequencing of Syn3A in exponential growth phase shows that Syn3A has an ori:ter ratio of 1.21. The dip at 22 kbp is likely due to natural evolution that resulted in deletion of the *tetM* gene. (C) Fluorescence imaging of JCVI-syn3B with a second copy of FtsZ labeled with mCherry (red), DNA stained with Hoechst 33342 (blue), and membrane stained with FM1-43FX (green). (D) Pie chart showing fraction of the nspected total 1,319 cells, that were annotated as prolate, dividing, and budding. (E) Visualization of simulated membrane (green 10 nm cubes), DNA (blue 10 bp particles), and FtsZ (red particles) at 70 and 95 minutes (top, bottom respectively). The simulated division morphology was generated geometrically based on two overlapping spheres.

Through fluorescence imaging of JCVI-syn3B, a variant of Syn3A with a “landing pad” system to mediate genetic modification,^30^ we observe that the minimal cell undergoes roughly symmetric division. Dividing cell shapes most frequently exhibit prolate (early cell division state) and dumbbell-like (late cell division state) morphologies. A few representative cells at different stages of the cell cycle are shown in Figure 4C. The gene encoding labeled FtsZ was introduced as an additional copy rather than a substitute for the original to minimize disruption of the division machinery. The distribution of morphologies for 1,319 analyzed cells (Figure S2) is shown in Figure 4D. Spherical cells appear larger than previous cryo-ET,^21^ but this discrepancy is consistent with observations for similar organisms where cell sizes can vary by hundreds of nanometers depending on the imaging method and sample preparation.^31^ The fraction of prolate and dividing cells observed in the imaging data is lower than expected based on our simulated cell cycle in which division begins approximately 60 minutes into a 105-minute cycle, which corresponds to roughly 40% of cells dividing for asynchronous cell cycles. This discrepancy is likely because our model does not explicitly simulate the kinetics and mechanics of FtsZ polymerization during division.

In addition to spherical and symmetrically dividing cells, we observed cells with “budding” morphologies and “cell-in-cell” intracellular membrane structures. A full-field-of-view Z-stack movie with overlays around all segmented cells is visualized in Video S2. Budding structures do not indicate that division of Syn3A occurs through budding, as an extracellular membrane structure that contained ribosomes but no DNA was previously annotated in cryo-ET of Syn3A.^21^ In late-exponential phase Syn3A, roughly 15% of analyzed cells contained cell-in-cell structures, statistics shown in Figure S2. Unfortunately the causes of these budding and cell-in-cell structures are not known, but even with undisturbed machinery to control division, irregular morphologies including cell-in-cell structures have been observed in other imaging of Syn3A.^19,32^

We combine the new fluorescence imaging and previous cryo-ET observations^21^ into two constraints on the morphology during our simulations such that the cell: 1) grows spherically from 200 to 250 nm in radius (~98% increase in cell volume) given our coarse-grained resolution of 10 nm lattice cubes and then 2) maintains a constant volume as the surface area grows during symmetric division until the end of the cell cycle. The dividing cell morphology is generated using a geometric approach where we treat the shape as two overlapping spheres that have the total instantaneous surface area and volume of the dividing cell. Although this does not include the kinetics and mechanics of FtsZ polymerization, the rate of change for the morphology is a direct result of the chemical reactions synthesizing the membrane components. An example of one simulated cell at two time points during division is shown for comparison to the experimental images. We visualize FtsZ for direct comparison to the fluorescent imaging and to show that in the absence of any implemented polymerization reactions, it does not localize to the division plane in our simulations. As shown in Figure S3, we tested a physics-based approach to generate dividing cell shapes using a discretized version of the Helfrich Hamiltonian implemented in the software FreeDTS.^33^ This method generates membrane shapes by minimizing constraints on the curvature and surface area to volume ratio obtained from our kinetic model. However, in the absence of a kinetic model of FtsZ polymerization to constrict the membrane at the division plane, the resulting morpholologies are more elongated than the shapes observed in the fluorescence imaging.

In our simulations, the predicted DNA replication and membrane doubling times correspond to an average B period of roughly 5 minutes (ori:ter - 1:1), C period 46 minutes (2:1), and D period 54 minutes (2:2). By uniformly sampling time points throughout the average simulated cell cycle, our simulations exhibit an ori:ter ratio of 1.28. In whole-genome sequencing (WGS) we calculate an average ori:ter ratio of 1.21 for exponential phase cells (Figure 4) and 1.0 for stationary phase cells (Figure S2). The consistency between predicted and experimental ori:ter ratios supports the assumption that Syn3A typically only initiates replication once per cell cycle. This agreement also suggests that our model recovers not only a realistic doubling time, but also realistic B, C, and D periods. As additional validation for the ori:ter ratio of 1.21, we analyzed DNA sequencing coverage data from a recent study that tracked the evolution of Syn3A over ~400 generations and found values ranging from 1.0 to 1.2.^20^

Additionally, our sequencing data revealed a significant drop in coverage at the *tetM* locus. *tetM* encodes the tetracycline resistance protein TetM, which was not needed in our growth conditions since tetracycline was absent from the medium. A similar observation was made in the evolutionary study mentioned above where mutations frequently arose in *tetM*.^20^ The authors suggested that the strong *tetM* promoter,^20^ combined with the absence of tetracycline, made *tetM* expression energetically costly,^34^ leading to frequent mutations that downregulated or disrupted it. While our data show a notable reduction in coverage rather than a complete deletion, this finding aligns with the idea that *tetM* is nonessential under these conditions.

### Formation of macromolecular complexes

A significant limitation of the previous static-structure and early cell cycle 4D simulations was that the number of ribosomes, RNAP, and degradosomes were constant and ribosomes and degradosomes did not diffuse.^1^ The assembly reactions and diffusion rules for these complexes are now simulated and are described in the Methods. The total counts of ribosomes, RNAP, and degradosomes over the course of a cell cycle are shown in Figures 5A-C. The counts of RNAP and degradosomes rapidly increase from 0 to 93 and 120 respectively because we initialize their components unassembled and they rapidly assemble in the first second of biological time. On average, the components are synthesized and assembled at rates such that the numbers of complexes come slightly short of doubling in an average cell cycle. This is due to the slight underproduction of proteins by our model, which is discussed in more detail below. We track each state of assembly for the complexes, so we can count the number of ribosomes that sit in incomplete states as shown in Figure 5. We find that on average, there are roughly 100 SSU and LSU complexes or intermediates that are waiting on ribosomal proteins to be synthesized to finish assembly as shown in Figure 5D.

**Figure 5:**
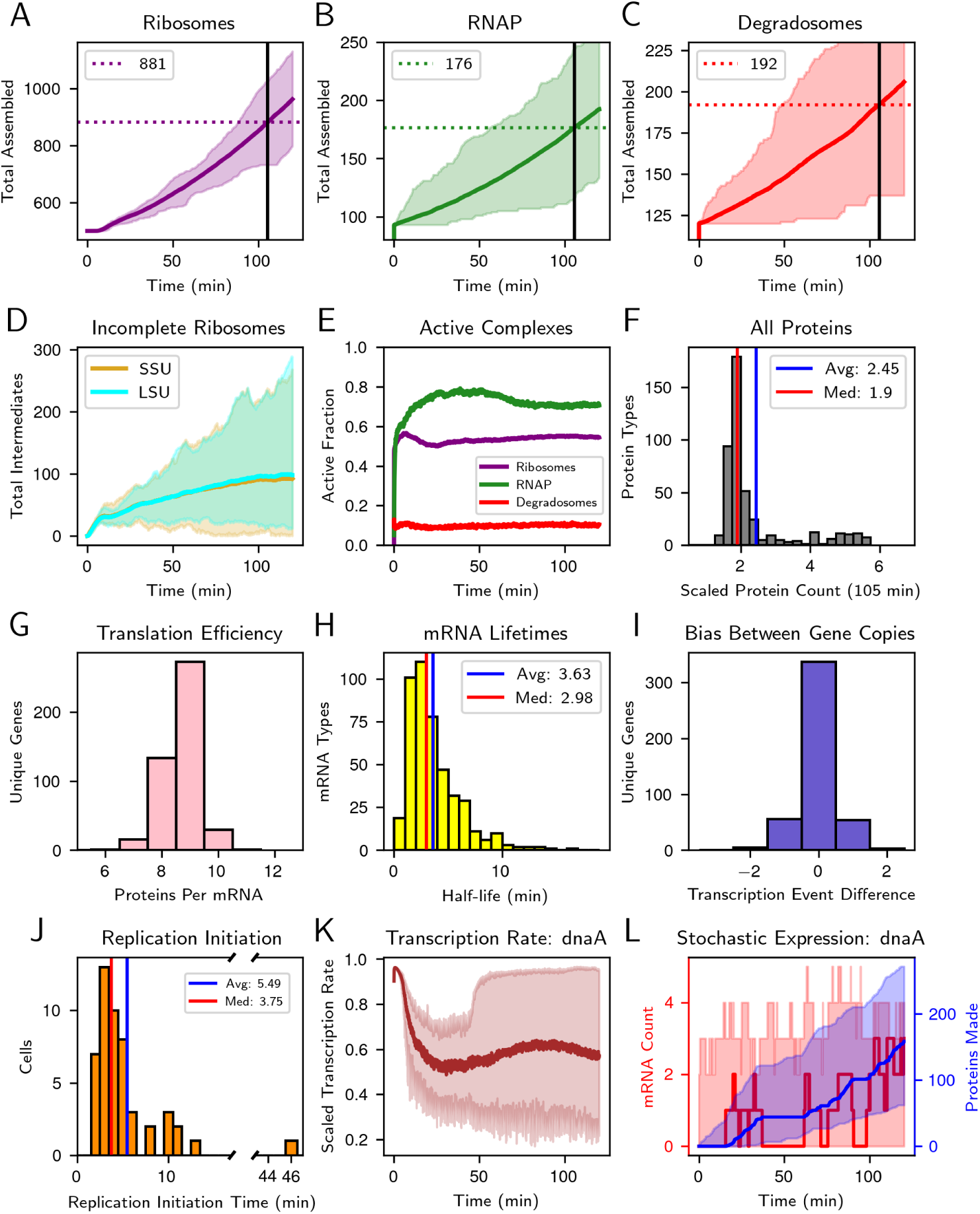
Dynamics of gene expression and macromolecular assembly. Cells have averages of 881 ribosomes (A), 176 RNAP (B), and 192 degradosomes (C) at the average predicted division time (105 min, vertical lines). (D) After a cell cycle, there are averages of roughly 100 incomplete large and small ribosomal subunits waiting for ribosomal proteins to be translated. (E) On average, roughly 55% of ribosomes are actively translating, 70% of RNAP are actively transcribing, and 10% of degradosomes are actively degrading at any one time. (F) Replication typically initiates around 5 minutes, but can initiate anywhere from 2 to 46 minutes. (G) Once a gene is duplicated through DNA replication, we measure the difference in number of transcription events between genes (transcription events gene copy 1 - gene copy 2). A positive value means the original gene copy is transcribed more frequently after replication, and a negative value corresponds to more transcription on the daughter. (H) Average number of proteins made per mRNA for all 452 genes. (I) Distribution of mRNA half-lives. (J) Scaled protein count (counts at 105 min divided by initial condition) show the model comes slightly short of doubling the counts of proteins in a cell cycle. (K) Scaled transcription rate of the *dnaA* gene (instantaneous rate divided by maximum possible elongation rate). (L) mRNA count and number of proteins translated for *dnaA* where the solid lines show the trajectory of a representative cell. All shaded regions in time-dependent plots represents the full range of the simulated population. Values for genomewide statistics are provided in Table S3.

### Activity of gene expression complexes and competition for mRNA

The activity of gene expression complexes (RNAP, ribosomes, and degradosomes) are quantities that are strongly tied to growth rate.^35^ For the conditions and parameters in our model, we predict that on average, roughly 55% of ribosomes, 70% of RNAP, and 10% of degradosomes are active at any one time throughout the cell cycle as shown in Figure 5E. The active fraction of degradosomes is lower than we previously predicted,^1^ but there is currently no available experimental comparison. The effects reducing the degradosome active fraction will be described in more detail below. The RNAP fraction is significantly higher than previously predicted. The higher active RNAP fraction is because our model now uses more informed assembly and stoichiometry to assemble RNAP complexes, which reduced the total count of complexes from 187 to 93 based on proteomics of the alpha subunit. Additionally, we found that the overall cell state and protein production is sensitive to the binding rate of the RNAP to promoters. This was already known because scaling the promoter strengths by an order of magnitude or less has been shown to affect the overall protein production in our previous models.^1,36^ The rate of RNAP binding to promoters presents itself as a global variable that is sensitive in the spatially-heterogeneous reactions. We do not treat co-transcription of polycistronic operons in these simulations, each gene is transcribed independently. However, we allow multiple RNAP to read the genes coding for rRNA, up to 7 per 23S gene and 4 per 16S gene based on an RNAP spacing of 400 bp (a conservative estimate from transcription measurements in *E. coli* ^35,37^).

The fraction of active ribosomes is higher than we predicted in our previous static-structure 4D model that used a ribosome distribution from cryo-ET,^1^ but the active fraction of ribosomes has been estimated to be as high at 85% in *E. coli*.^35^ In giving the ribosomes the ability to diffuse in these simulations, we had to sacrifice a critical effect in the overall translation dynamics: polysomes. The fraction of ribosomes involved in polysomes has been measured or estimated in several bacteria including: up to 70% in *E. coli*,^38^ up to 26.2% in *Mycoplasma pneumoniae*,^39,40^ and 20 to 40% in Syn3A.^21^ Given the fractions of ribosomes in polysomes observed in experiments, our predicted fraction of active ribosomes is not unreasonable in the absence of polysomes. We do not include polysomes because each ribosome is independent of all other ribosomes in the RDME representation.

One of our key metrics in evaluating the accuracy of our models has been the ability to double the counts of proteins over the course of a cell cycle.^1,36^ We again show this metric here in Figure 5F and the average number of every protein in the simulation are provided in Table S3. Our distribution is not as narrow as it was in the well-stirred model, now exhibiting a tail out to almost 6 times the proteomics counts for some proteins. This tail is composed mostly of proteins with initial counts of 10 particles (10 or fewer measured in proteomics^1,36^) and the rest have initial counts of less than 50. However, the median of the distribution falling below 2 shows that most proteins are underproduced in our simulations. We observe several underproduced proteins whose counts are only increased to 1.25 to 1.5 times their initial counts after a cell cycle, all of which have long protein sequences (gene length *>*3 kb). We found three key effects come into play in the rate of protein production in Syn3A: translation efficiency, mRNA lifetimes, and transcription rate. Our simulations lack the translational effects of polysomes, meaning each mRNA can only be translated by a single ribosome at a time. We showed previously that polysomes play a significant role in doubling the number of proteins whose mRNA can be translated by more than one ribosome simultaneously.^1^ In the absence of polysomes, proteins with longer sequences that would benefit from simultaneous translation from multiple ribosomes are underproduced.

Because we track every transcription, translation, and mRNA degradation event, we can calculate the average translation efficiency and mRNA half-life for every type of mRNA in Syn3A as shown in Figures 5G and H, respectively. The previous static-structure 4D simulations exhibited average and median half-lives of 1.97 and 1.48 minutes, respectively.^1^ The most impactful change to these values was an adjustment of the ratio of the binding rates for mRNA to ribosomes and degradosomes. Using the binding rates of mRNA to ribosomes and degradosomes from the previous model, preliminary simulations showed that the simulated cells would fall significantly short of doubling the counts of most proteins. We found that the ratio of these binding rates is among the most sensitive in the 3D stochastic kinetics. The final parameter set increases the binding rate of mRNA to ribosomes by a factor of 1.3 (30% increase) and reduces the binding rate of mRNA to degradosomes by a factor of 0.3 (70% decrease). While these seem like large changes, both represent changes of less than an order of magnitude, which is relatively small in the context of cell-wide kinetics. These changes are reflected in the distribution of mRNA half-lives and translation efficiencies showing that mRNA are longer lived and more proteins can be translated per mRNA during the longer lifetimes. The increased mRNA lifetimes and translation efficiency then give us a distribution of protein production that comes closer to doubling the counts of most proteins. There are more values for parameters that could be tested, but testing multiple parameter sets in these simulations is computationally expensive, and other biological factors like polysomes should be integrated before further varying these sensitive parameters.

We analyzed the difference in transcription events per cell cycle between the both copies of each gene on the simulated daughter chromosomes (shown in Figure 5I). Most gene copies are unbiased, and we found no clear trend in the identities of genes that exhibit bias toward a specific gene copy. However, we found 81 genes that were not transcribed in one to three cell cycles among the 50 simulated cells. This is not an unreasonable prediction because some proteins perform their functions infrequently. As long as some proteins are present, the cell can live, therefore gene expression does not need to happen every cell cycle. The number of cells in which each gene goes untranscribed is listed in Table S3. The genes that go untranscribed are all low in proteomics value (*<*60) and the majority of them are genes of unknown function.

### Replication initiation kinetics are sensitive in 3D

DnaA controls the timing of DNA replication initiation through polymerization reactions that unwind DNA near the origin. We use a model based on the kinetics from our previous well-stirred simulations.^1,36^ In short, we treat the origin of replication as an individual particle on the RDME lattice that follows the origin monomer on the BD DNA. The RDME particle undergoes state changes as more DnaA bind and the simulated kinetics are described in Methods. We previously used the reported average on and off rates of 100 mM^−1^s^−1^ and 0.55 s^−1^ for DnaA domain III binding to ssDNA in our well-stirred kinetic model.^1,36^ However, when we implemented these rates in the 4D model, replication did not initiate within the first 60 minutes of the cell cycle. Due to the computational expense of these simulations, we did not explore this effect in depth and increased the strength of DnaA binding to single-stranded DNA by increasing the on rate to 140 mM^−1^s^−1^ and reducing the off rate to 0.42 s^−1^. Both pairs of on and off rates were measured experimentally in the same *in vitro* single-molecule FRET study.^41^ 100 mM^−1^s^−1^ and 0.55 s^−1^ correspond to the average measured rates. 140 mM^−1^s^−1^ and 0.42 s^−1^ correspond to rates measured for a DNA structure similar to the origin of Syn3A consisting of both an AT-rich ssDNA and a dsDNA sequence containing two DnaA domain IV binding sites. Even though the differences between rates are less than 50% each, replication initiates in almost every simulated cell within the first 15 minutes of the cell cycle and the range of filament lengths at the time of replication initiation range from 20 to 23 DnaA. Distribution of replication initiation times is shown in Figure 5J. Even the cell that did not initiate replication until 46 minutes into its cell cycle successfully completed replication and division within 2 hours.

### Genetic information reactions and metabolism are co-dependent

Although we will not discuss the dynamics of metabolism thoroughly here, it is still a critical factor driving the overall progression of the cell cycle in the 4DWCM. The pools of metabolites dictate the synthesis rate of macromolecules like the DNA and RNA and the uptake rates of lipids determines the rate of growth for the membrane. We show the concentrations of all metabolites and fluxes through all metabolic reactions in Data S1. Additionally, the energetic costs of all ATP-activated processes were once again quantified and are shown in Figure S4, but we do not observe any significant differences in ATP costs from the previous well-stirred model. As an example of the connection between metabolism and genetic information, we show the transcription rate for the *dnaA* gene in Figure 5K. The average transcription rate is reduced to roughly 55% of the maximum transcription elongation rate (corresponding to roughly 11 nt/s) with a range anywhere from roughly 30 to 80% until the volume has doubled. Once the volume doubles, some cells fully recover a maximum transcription elongation rate and some are still slowed down. Variations in transcription speed are due to fluctuations in the pools of NTPs, as the elongation rate depends directly on the instantaneous NTP concentrations as described in the Methods. A significant difference from the previous well-stirred model is that this model does not accumulate large quantities of NTPs due to re-balancing of the nucleoside uptake rates, the glucose uptake rate, and the glycolysis reaction fructose 1,6 bisphosphate aldolase (FBA). The molecules that are most commonly limiting the rate of transcription are UTP and GTP, both of which have average pools around 0.3 mM but are sometimes fully depleted. Concentrations of 1 to 10 mM GTP and 8 mM UTP have been estimated for *E. coli*,^42^ but no measurement has been performed yet in Syn3A. While the NTP pools here seem too low, the uptake rates are generating excess pools of NTP precursors like uridine and adenosine. Additionally, there are excess pools of all dNTPs as can be seen in Data S1. Therefore, we are likely missing some balancing effects between NTP and dNTP pools in the metabolic rates, an example of which is metabolic inhibition. The average transcription rate being reduced is directly connected to protein formation by affecting the overall number of transcripts that are made. While the average transcription elongation rate makes transcription look like a continuous process, we show that the mRNA population still depends on the stochastic probability of transcribing the gene and degrading the mRNA through the mRNA counts of *dnaA* for a single cell shown in Figure 5L. We also show the total number of DnaA proteins made as a function of time for the same cell to show the bursty nature of gene expression; proteins can only be made when mRNA are present.

### Partitioning of macromolecules to daughter cells is stochastic

Because we treat all proteins, RNA, DNA, and selected complexes in a 3D representation, our model automatically predicts the partitioning of these molecules to daughter cells once the division model isolates the daughter cells’ cytoplasms and separates their membranes. The random diffusion of particles on the lattice determines their locations at the time of division. We show a representative selection of particles and how their particles were partitioned in Figure 6: ribosomes, degradosomes (a peripheral membrane particle), PtsG (a transmembrane protein), and GapDH (a monomeric cytoplasmic protein). The population distributions appear to be random distributions approaching a binomial distribution. For each particle type chosen to select its region, the cell-to-cell variations show that our simulations have no significant bias toward one daughter cell over another for partitioning particles after division. Of note, the distribution for degradosome count differences shows peaks at the edges of the distributions. This is because of the DNA coarse-graining on the RDME lattice. If large portions of the DNA are pushed into contact with the membrane, the DNA lattice sites exclude peripheral membrane proteins like degradosomes, resulting in a bias against that side of the dividing cell.

**Figure 6:**
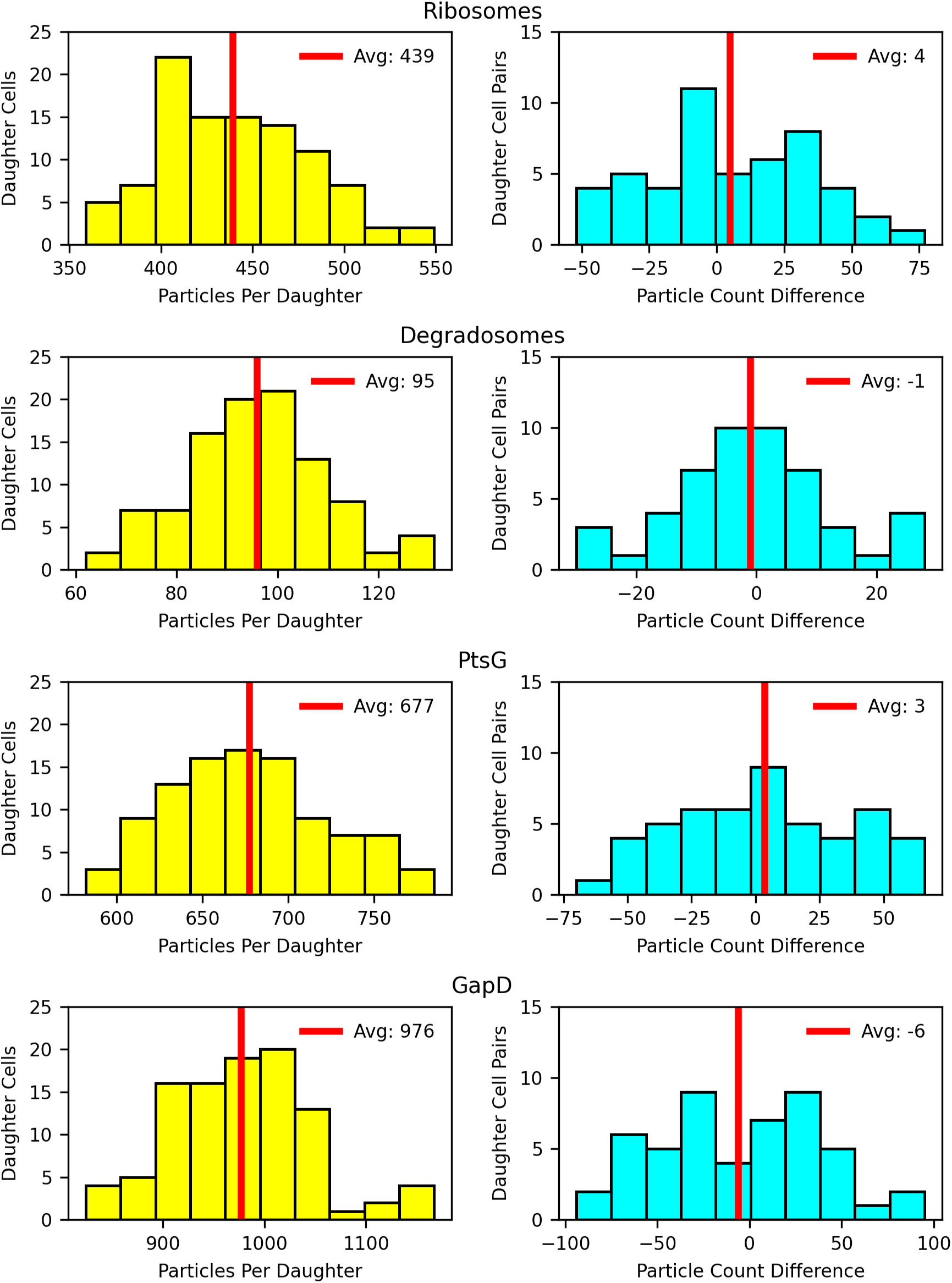
Partitioning of macromolecules to daughter cells resulting from random diffusion on the RDME lattice after division. Left: Distribution of counts per cell among the population of divided cells. Right: Difference in particle counts between pairs of daughter cells. Positive values correspond to more particles in the daughter cell in the positive z direction (relative to the division plane) and negative values correspond to more particles in the daughter cell in the negative z direction. All counts were taken from the 105 minute time point corresponding to the average predicted division time.

## DISCUSSION

Cells are not well-stirred reactor systems. Their intracellular environment is spatially heterogeneous and consists of many slow-moving, low-population components that need to encounter each other in 3D space for them to react and perform their biological functions. Achieving spatial reaction-diffusion dynamics has been a significant challenge in the field of whole-cell modeling. Here, we presented a 4DWCM over an entire cell cycle of the genetically minimal cell, JCVI-syn3A. Simulating an entire cell cycle in 3D posed several challenges, the most significant including: making chromosomes dynamic and partitioning them to daughter cells, updating morphology and connecting it to the hybrid stochastic-deterministic chemistry, and moving ribosomes. Overcoming each of these challenges required advances and integration of new computational methods and algorithms. By leveraging the capacity of our GPU resources, we separated BD simulations of the chromosome(s) from the LM RDME simulation onto two individual GPUs. Parallelization of these two tasks dramatically sped up the time to simulate a cell cycle from over two weeks down to 4 to 6 days.

We found that DNA replication initiation and competition between translation and mRNA degradation are strongly sensitive to spatial heterogeneity. The second-order binding rates that characterize the dynamics of these processes were found to be sensitive in affecting the overall progression of the cell cycle. For example, a slight reduction in affinity of DnaA to form the filament near the origin resulted in replication never initiating in testing. A slight reduction in the affinity of mRNA to bind to degradosomes relative to binding to ribosomes resulted in protein production that comes closer to doubling protein counts in a cell cycle.

Among the greatest advantages of our 4DWCM is the ability to predict many measurable quantities at the same time, the equivalent of thousands of simultaneous experiments. We predict a doubling time of 105 minutes based on surface area growth the depends on lipid and membrane protein synthesis rates, which agrees very closely with the experimentally measured doubling time of 105 minutes.^18^ Additionally, our combined B, C, and D periods with a single replication event per cell cycle result in an average ori:ter ratio of 1.28, which agrees well with the ratio of 1.21 measured in DNA sequencing. Many quantities are still predictions for Syn3A that have yet to be experimentally validated, for example the genome-wide activity statistics for gene expression complexes. We have made comparisons to other bacteria where data is available, however there can be large variations from organism to organism in the behavior exhibited by individual processes. For example, the average mRNA half-life can vary significantly from organism to organism.^43–45^

### Limitations of the study

Among the most significant limitations of our 4DWCM is the required computational time and resources to achieve statistically significant sampling. As mentioned above, our simulations require 4 to 6 days of computational time of two high-performance computing GPUs per cell cycle. To obtain the 50 replicate cell cycles presented here required roughly 15,000 GPU hours on NVIDIA A100 GPUs. This limits the sampling that can be performed to small populations of cells. Despite this long compute time, and although the coarse-graining of our model means that we lack the physical accuracy of the predicted interactions from methods like molecular dynamics, we simulate hours of biological time. Well-stirred simulations can take as little as a few hours per cell cycle and can simulate multiple cells in parallel.^1,4,11,36^ We previously showed that 4D simulations can be used to better constrain and parameterize related well-stirred models. While we cannot simulate large numbers of cells with 4D models and well-stirred models lack the predictive power of 4D models, we can build the crossroads between the two levels of complexity to accelerate the development of more predictive whole-cell models.

Another computational limitation comes with the frequency at which frames of the RDME simulation are written to the trajectory file. We previously hypothesized that the lifetime of a mRNA depends on the spatial proximity of the gene to the membrane where the degradosomes are located.^1^ The proximity of the *fbaA*/0131 gene particle on the RDME lattice for the mother chromosome and both daughter chromosomes to the nearest membrane lattice site in a single cell is shown in Figure S3. Gene-membrane proximity was averaged over the entire cell cycle and we tried to correlate the proximity to bulk properties including mRNA degradation rate, mRNA half-life, and protein production rate. We found no statistically significant correlations. An example of the correlation to single-cell mRNA half-lives for *fbaA* is shown in Figure S3. To better quantify the relation between spatial proximity to the membrane and mRNA lifetime would require us to record every frame of the RDME trajectory. Currently, we record one frame every 1 second of biological time. In the implementation of the 4DWCM every mRNA is indistinguishable from other mRNA of the same type. Therefore, unless we follow its diffusion every timestep, we cannot explicitly track fate of an individual mRNA particle. Unfortunately, recording every timestep of the RDME trajectory is currently prohibitive as we estimate that a single trajectory file would require more than 80 TB of storage.

Although our model has made many significant predictions, it is still far from being complete and there are some important ingredients yet to be included: transcription of polycistronic RNA from operonal structures, cooperative genetic information processing reactions, a kinetic model of FtsZ to drive cell division, assembly reactions for all macromolecular complexes, and a defined biological system that partitions the chromosomes to daughter cells. The assumptions of the current transcription model have two known inaccuracies: independent transcription for all genes and promoter strengths assigned based on proteomics counts, both discussed in the Methods. It is known that many genes are transcribed cooperatively into polycistronic RNA,^20^ but this will require significant complexification of the reaction model and tracking of transcriptional states. Our current and previous models have all parameterized the promoter strengths based on the quantitative proteomics data,^1,18,36^ but once the transcription model is more complete it will be important to instead constrain expression levels based on quantitative transcriptomics. Incorporation of polysomes will require integration of another simulation method to handle the diffusion of ribosomes separately. Other cooperative processes like coupled transcription-translation and membrane protein translation-translocation are emerging as important factors in gene expression.^39,40,46^ A well-stirred model of assembly reactions for all complexes is in development, and this model will need to be adapted to 4D with the incorporation of coupled translation-translocation of membrane proteins. While polymerization of FtsZ has been studied, the dynamics have not been characterized in Syn3A. A realistic model of division including FtsZ dynamics will require hybridization of a kinetic model of polymerization with a physical model that generates realistic division morphologies from the FtsZ kinetics.

Finally, there are several simplifications of the current model of chromosome partitioning to daughter cells. Partitioning of chromosomes was handled using a weak fictitious repulsive force between the daughter chromosomes. Mechanisms that are known to partition chromosomes in other organisms such as the Min system in *E. coli*,^47^ or the ParAB*S* system in *B. subtilis* or *C. crescentus* ^48^ have either been removed from or are not annotated in Syn3A. Unfortunately, we cannot simulate enough Brownian dynamics time steps to determine if entropic segregation is sufficient to partition the chromosomes to daughter cells. While entropic segregation is one hypothesis, we made several simplifying assumptions about SMC behavior in our model that could be refined and complexified to better partition the chromosomes.^49–52^ Additionally, we assumed a fixed dwell time of 4 second and assume that all SMCs remain bound during that 4 second dwell time. We have started exploring procedures with varied SMC physics and dwell times, and a preliminary study as shown in Video S3 exhibits chromosome configurations suggesting that it is likely possible that our fictitious force could be replaced by a better SMC model. However, the procedures that could partition chromosomes with SMC are currently too computationally expensive.

## Supporting information

Key Resources Table

## RESOURCE AVAILABILITY

### Lead contact

Requests for further information and resources should be directed to and will be fulfilled by the lead contact, Zaida Luthey-Schulten.

### Materials availability

This study did not generate new materials.

### Data and code availability

- All original code to run and analyze the 4DWCM will be made publicly available on Github and Zenodo upon acceptance of publication.
- Files containing all particle counts for the 50 simulated cells will be made publicly available on Zenodo upon acceptance of publication. Four representative LM RDME trajectory files from the 4DWCM are also provided.
- DNA sequencing data has been deposited in the NCBI SRA under PRJNA1257452.
- Any additional data reported in this paper and the information required to reanalyze the data is available from the lead contact upon request.

## ACKNOWLEDGMENTS

T.W., E.F., J.K., J.I.G., and Z.L.S. are supported in part by NSF MCB 2221237. Z.L.S., T.H., and A.Me. are supported in part by the NSF Science and Technology Center for Quantitative Cell Biology (NSF DBI 2243257). W.P. research is supported by the Novo Nordisk Foundation (grant No. NNF18SA0035142 and NNF22OC0079182) and Independent Research Fund Denmark (grant No. 10.46540/2064-00032B). Z.R.T.: Research reported in this publication was supported by the Cancer Center at Illinois - Beckman Institute Postdoctoral Fellows Program sponsored by the Cancer Center at Illinois and the Beckman Institute for Advanced Science and Technology, University of Illinois Urbana-Champaign. The content is solely the responsibility of the authors and does not necessarily represent the official views of the program sponsors.

The authors would like to thank Jay Cournoyer for assistance in culturing the JCVI-synX cell lines. We thank Glenn Fried for assisting with access to the shared facilities at the Carl R. Woese Institute for Genomic Biology at UIUC. We thank Chris Fields, Alvaro Hernandez, and Chris Wright from the Roy J. Carver Biotechnology Center sequencing facility at UIUC for assistance with collecting and processing the DNA sequencing data. The authors also thank Martin Gruebele, Rohit Bhargava, Kim S. Wise, Clyde A. Hutchison III, and Hamilton O. Smith for helpful discussions.

## AUTHOR CONTRIBUTIONS

Conceptualization, Z.R.T and Z.L.S.; methodology, Z.R.T., A.Ma., T.A.B., and B.R.G.; software, W.P., T.W., and H.L.; investigation, J.K., E.F., J.Q, Y.G., T.L., L.S.; writing-original draft, Z.R.T, A.Ma., and Z.L.S.; funding acquisition, Z.L.S.; resources, A.Me. and J.I.G.; supervision, A.Me., T.H., J.I.G., and Z.L.S.; writing – review & editing, all authors.

## DECLARATION OF INTERESTS

The authors declare no competing interests.

## DECLARATION OF GENERATIVE AI AND AI-ASSISTED TECHNOLOGIES

During the preparation of this work, the author(s) used ChatGPT in order to write Python functions to generate more aesthetically appealing plots. After using this tool, the author(s) reviewed and edited the content as needed and take(s) full responsibility for the content of the publication.

## SUPPLEMENTAL INFORMATION INDEX

Figures S1-S4 and their legends are provided in a supplemental PDF.

Table S1. Initial conditions and average concentrations for all intracellular metabolites. Concentrations of nutrients in the simulation growth medium. (Excel file)

Table S2. Complete set of kinetic parameters used for metabolic reactions. Free energy of central, nucleotide, and lipid metabolic reactions including standard and reaction free energies. Average simulated fluxes for all metabolic reactions. (Excel file)

Table S3. Genome-wide numeric values for statistics of genetic information processes and average particle counts for related molecules. (Excel file)

Data S1. Trajectories of all time-dependent metabolite concentrations and metabolic fluxes. Solid lines represent the population averages. Shaded regions represent the ranges observed for the entire population of simulated cells. (PDF file)

Video S1. Video of the cell cycle of a simulated cell. The visualization includes the Brownian dynamics chromosomes, lattice membrane, ribosomes, and transmembrane sugar transporter PtsG.

Video S2. Full-field view of a subpopulation of fluorescent labeled JCVI-syn3B-FtsZ:mCherry cells. Color scheme is the same as Figure 4C. Scan goes through roughly 2 microns of Z-stack images.

Video S3. Video showing preliminary study of chromosome partitioning by the action of SMC alone. Green is the mother chromosome, pink and blue are the daughter chromosomes.

## STAR METHODS

### Key resources table

The KRT is provided in an independent Word document.

### Fluorescence Imaging of Dividing Cell Shapes

#### Growth Medium

SP4 part 1: Resuspend 3.5 g Mycoplasma Broth Base, 10 g Bacto Tryptone, and 5.3 g Bacto Peptone in distilled water to a final volume of 600 mL. Adjust pH to 7.5. Autoclave for 15 minutes at 121*^◦^*C.

SP4 part 2: Combine the following: 25 mL Glucose (20% w/v stock), 50 mL CMRL 1066 (10X stock, w/o phenol red, w/o bicarbonate, w/o Gln), 6 mL Sodium Bicarbonate (7.5% w/v), 5 mL L-glutamine (200 mM), 35 mL Yeast Extract Solution (15% w/v), 100 mL TC Yeastolate (2% w/v), 170 mL KnockOut Serum Replacement, 2.5 mL Penicillin G (400,000 U/mL), and 1.5 mL Phenol Red (1% w/v). Filter sterilize (0.2 *µ*m) and store at 4*^◦^*C.

SP4 medium: Combine SP4 part 1 and SP4 part 2 with a 1.5:1 ratio under sterile conditions.

#### Genetic Labelling of FtsZ in JCVI-syn3B

JCVI-syn3B is a genetic variant of Syn3A with a “landing pad” system to mediate genetic modification that has the same doubling time as Syn3A when no genetic modifications have been inserted in the landing pad. Genetic insertion of Ftsz-mCherry (JCVISYN3A 0522 fused with an *mCherry* gene) was performed as described previously for insertion of a single gene to the Syn3B genetic landing pad.^30^ To minimize disruption to the cell division machinery, the original gene copy was left intact and mCherry was added at the C-terminal end of the genetically inserted second copy of FtsZ.

#### Cell Culture and Sample Preparation

Minimal JCVI-syn3B cells expressing the FtsZ-mCherry fusion protein were cultured in SP4 medium supplemented with KnockOut Serum Replacement (ThermoFisher #10828028). For each experiment, cells were freshly prepared from frozen stocks. A small portion of the frozen cells was scraped and transferred to 20 mL of culture medium, followed by static incubation at 37*^◦^*C for 24 h.

Cells were fixed during the logarithmic growth phase by adding 2 mL of 32% paraformaldehyde (Electron Microscopy Sciences #15680) and incubating for 30 min on a nutator. Fixed cells were then harvested by centrifugation at 4,500 × g for 10 min. The resulting pellet was resuspended in 1 mL of phosphate-buffered silane (Corning #21-040-CV) and seeded onto an 8-well chambered coverslip (Fisher Scientific, #12-565-470) coated with poly-D-lysine (Gibco, #A3890401).

After approximately 6 hours of incubation, cells were rinsed twice with PBS and sequentially stained with 1 *µ*g/mL Hoechst 33342 (Thermo Scientific, #62249) for 10 min at room temperature, followed by pre-cooled 5 *µ*g/mL FM1-43FX (Invitrogen, #F35355) for 1 min on ice. After each staining step, cells were washed three times with PBS for 5 min to remove excess dye.

#### Airyscan Imaging

A Zeiss LSM 900 confocal microscope equipped with an Airyscan 2 module was used to acquire three-color images of fixed JCVI-syn3B FtsZ-mCherry cells. All images were captured in Airyscan mode with an xy pixel size of 35-nm and a z-step of 130-nm for 3D imaging. Fluorescence signals were collected for 1.1 *µ*s per pixel per color channel. Sub-diffraction-limited image processing was performed using ZEN software (Zeiss).

The acquired 3D images were first manually segmented to identify single, dividing, and budding cells (a schematic of the image analysis workflow has been prepared). Cells exhibiting stable fluorescence intensities across the z-stacks were selected for further analysis (Video S2: whole-view image with segmentation boxes). Syn3A/B cells do not have machinery for cell movement (e.g. flagella), causing the cells form clusters as the population grows. Clustering of the cells frequently prohibits classification. In total, we segmented 2,968 cells, but due to the weak adhesion of cells to the surface and clustering, approximately 1,649 cells were not suitable for analysis (Figure S2).

For each cell, a single z-plane showing the clearest membrane structure was typically selected for segmentation. When a single plane did not sufficiently capture the cell’s morphology, a maximum intensity projection of multiple z-planes was used instead.

Segmented single cells were further analyzed by fitting their membrane images to a Gaussian-distributed intensity function, where the intensity decays with increasing distance from the ellipsoidal edge. From the fitting results, the equivalent diameter and aspect ratio (minor axis length / major axis length) were calculated to characterize cell size and shape. Cells were classified as prolate if (1) their equivalent diameter exceeded 0.6 *µ*m and (2) their aspect ratio was below 0.8 (Figure S2).

### DNA Sequencing

#### Sample Preparation

A frozen stock of Syn3A-mCherry was provided by the JCVI. Frozen stocks were scrapped and inoculated into 10 mL of SP4 media (see formulation above) under BSL2 conditions and incubated at 37 *^◦^*C with shaking. Once the media acidifies (∼2 days), cells are used for experiments.

#### Growth Curve Experiments

48 hours post thaw, 1 mL of stationary phase Syn3A-mCherry culture was collected and centrifuged at 12,000 g for 10 minutes. The cell pellet was then resuspended in 1 mL PBS. 80 *µ*L of culture was used to seed 20 mL SP4 media cultures, one for each timepoint in triplicate. Cultures were left to grow at 37 *^◦^*C with rotation for 8 (exponential phase) or 21 hours (stationary phase). After growth, cultures were centrifuged at 13,000 × g for 10 minutes at 4 *^◦^*C. Cell pellets were snap-frozen and stored at −80 *^◦^*C until processed.

#### Genomic DNA extraction and Sequencing

Syn3A-mCherry frozen cell pellets were processed in parallel using the TrueLink gDNA extraction kit (ThermoFisher #K182001) as per manufacturer instructions. gDNA concentrations were quantified using Qubit, as per manufacturer instructions.

The shotgun genomic libraries were prepared with the Illumina DNA Prep kit (Illumina, #20060060). The library pool was quantitated by qPCR and sequenced on one MiSeq flowcell for 251 cycles from each end of the fragments using a MiSeq 500-cycle sequencing kit version 2. Fastq files were generated and demultiplexed with the bcl2fastq v2.20 Conversion Software (Illumina). Phix DNA is used as a spike-in control for MiSeq runs.

#### DNA coverage analysis

Sequencing quality was uniformly high for all replicates, with average Phred quality scores exceeding 35 across the majority of read positions, as reported by MultiQC. Raw sequencing reads were aligned to the reference genome using Bowtie 2 v2.5.4,^54^ and coverage profiles were generated using samtools v1.21.^55^ For plotting the coverages in Figure 4B and Figure S2F, we applied Gaussian smoothing to the data using a window with a standard deviation of 1000 bp generated by the scipy.signal.windows.gaussian function, and convolution was performed in the Fourier domain using the scipy.fft module.

For each coverage profile we fit a line across half of the genome from the terminus to origin (genome positions [271,690-543,379]). We decided not to include genome positions [0-271,689] in the fitting process because genomic coordinates [0-271,689] contained more dips in the data, such as the deletion of the *tetM* gene around [20,200-21,950], and furthermore replication should be roughly symmetric in both directions from the origin. The values of the fitted line at its end-points were taken as proxies for the origin and terminus coverage, respectively. For exponential growth phase, we obtained ori:ter ratios of 1.20 (replicate 1: 2120/1760), 1.21 (replicate 2: 2020/1670), and 1.21 (replicate 3: 1910/1580). In the stationary phase, the corresponding ratios were 0.98 (replicate 1: 1745/1785), 0.99 (replicate 2: 1594/1616), and 0.97 (replicate 3: 1583/1624).

We used the same fitting method for the coverage profiles generated from DNA sequence datasets of 10 evolutionary endpoints that were deposited alongside a recent study.^20^ The ratios were 1.06 (endpoint a1: 365/345), 1.10 (endpoint a2: 217/197), 1.05 (endpoint a3: 281/267), 1.13 (endpoint a4: 268/237), 1.12 (endpoint a5: 200/178), 1.07 (endpoint a6: 158/148), 1.02 (endpoint a7: 246/242), 1.12 (endpoint a8: 195/174), 1.19 (endpoint a9: 238/200), and 1.16 (endpoint a10: 243/210).

#### Construction of the Syn3A 4DWCM

The remainder of the methods describe the methods used in the 4DWCM. We describe the initial conditions, the simulation methods, the hybrid algorithm the connects the simulations methods, and the physical and chemical details for individual processes. We also include a discussion of the construction of the computational environments used to simulate the 4DWCM.

### Initial Conditions

#### Computational Growth Medium

In Thornburg et al., 2022, we published a defined growth medium composition for Syn3A in which the exact concentration of each nutrient was controlled except for lipids and fatty acids.^1^ We used this formulation to determine the concentration of extracellular nutrients in the simulations. Because we only simulate a single cell, we assume that the depletion of nutrients from the medium is negligible relative to bulk solution over the course of a single cell cycle. We therefore set the growth medium concentrations as fixed constants in the simulation. The exact concentrations used in the simulated growth medium are provided in Table S1.

#### Metabolites

We previously generated a set of initial concentrations for intracellular metabolites from scaling metabolomics values reported for *E. coli*.^1^ We did not modify these values from the previous models, and they are re-stated in Table S1.

#### Macromolecular Complexes

RNAP and degradosomes are both initialized with counts of zero for the number of assembled complexes. The individual protein subunits are randomly distributed throughout the cytoplasm and then quickly assemble early in the simulation. We find that the vast majority of RNAP and degradosome complexes that can form through the initial proteomics are assembled in *<*1 second of biological time in our simulations.

Each replicate cell is initialized with 500 ribosomes. The previously published cryo-electron tomograms of Syn3A showed that the ribosomes are uniformly distributed throughout the cytoplasm.^21^ Based on this observation, we randomly distribute the initial 500 ribosomes throughout the cytoplasm by randomly sampling a uniform distribution. Additionally, to account for excluded volume effects, no two ribosome center of mass particles are placed in the same lattice site. For each ribosome center of mass particle, we then build a cross consisting of 7 lattice sites around the center of the mass that are assigned the “ribo center” site type for the center of mass and “ribosomes” for each lattice site sharing a face with the center site (shown in Figure S1).

#### Proteins

All non-ribosomal proteins are initialized to their proteomics values reported in Thornburg et al., 2022.^1^ For cytoplasmic proteins, the particles are randomly distributed throughout the cytoplasm, DNA, and outer cytoplasm (membrane periphery) regions at ratios of 1/2, 1/3, and 1/6 roughly reflecting the ratio of the volumes of the three regions to achieve a uniform random distribution of proteins throughout the cell. The particles for all transmembrane proteins are randomly distributed throughout the membrane region. The peripheral membrane protein particles are randomly distributed throughout the outer cytoplasm region.

Ribosomal proteins are initialized with reduced counts from the reported proteomics values. We assume that the counts reported in the proteomics also include the proteins incorporated into ribosomes. To determine the number of free ribosomal proteins in the initial conditions, we subtracted the number that would be incorporated into ribosomes (500 per protein) from the respective protein’s proteomics count. However, not all ribosomal proteins have enough counts detected in the proteomics to form 500 ribosomes. We believe this is likely due to proteins stuck to rRNA that stay stuck during the trypsin digestion process, therefore reducing the number of proteins detected in mass spec. In this case, we assume that there are enough proteins to form 500 ribosomes and add an additional 25 proteins to the initial condition for the respective ribosomal protein. We chose 25 proteins because it has been reported that roughly 5% of ribosomal proteins are not ribosome-associated per ribosomal protein.^56^

The initial counts for the number of free proteins for the entire proteome are provided in the Supplemental Information.

#### RNA

All rRNA are assumed to be in complete ribosomes at the start of the simulation, so the number of all free rRNA are initialized to 0. We use the same initial conditions as the previous model for counts of tRNA. We distribute a total of 200 tRNA per isoform for the 20 types of tRNA. This was chosen by scaling the total number of tRNA observed in *E. coli* to the same concentration in Syn3A, which corresponds to a total of roughly 6,000 tRNA molecules per cell for a 200 nm radius Syn3A cell.^1^ Syn3A has a total of 29 tRNA isoforms among the 20 types of tRNA.

In the absence of quantitative transcriptomics, we used the average counts of mRNA from our previous model as the initial conditions. Each cell is initialized with a different count of each mRNA. To determine the initial count for each type of mRNA, we randomly sample a Poisson distribution whose average value corresponds to twice the average count observed in the previous model.^1^ We sample distributions with twice the average because in initial testing, we were already able to observe the average total mRNA content of cells with updated kinetic parameters for genetic information processing. When we quantified the average total mRNA count using the updated parameters, we observed the total mRNA count to be double the total observed in the previous model (400 vs 200 total mRNA). We make the assumption that the individual mRNA abundances are also roughly twice the average counts observed in the previous model. The general increase in mRNA abundance is a result of increasing mRNA stability (and therefore lifetimes) by increasing the binding rate to ribosomes and decreasing their binding rate to degradosomes. The binding rates are discussed below and the increased lifetimes are shown in Figure 5. A complete table of the average counts used to sample the Poisson distributions are provided in the Supplemental Information.

#### Chromosome Configuration

The initial chromosome configuration is generated using the program sc chain generation. This program iteratively grows the circular chromosome into a spherical boundary as a series of spherocylinders representing a set number of beads. The exact algorithms are described in depth by Gilbert and coworkers.^6^ We chose to iteratively generate the chromosome over 4 stages: 2000 12-bead, 8000 6-bead, 18000 3-bead, and finally 54338 1-bead growth stages. The final stage corresponds to 10 bp per bead for the 543 kbp chromosome of Syn3A. We generate this configuration within a spherical boundary 1900 Å in radius, a reduced volume from the full cell to ensure that the chromsome does not intersect with membrane lattice cubes in the RDME representation.

Once the initial configuration has been generated, we impose the chromsome configuration on the RDME lattice. To do this, we iterate over each of the 54338 beads in the chromosome and divide their x, y, and z coordinates by the resolution of the RDME lattice (10 nm). Then, for each coordinate, we set the corresponding lattice site to the DNA site type. Some particles may correspond to the same lattice site, in which case the site is still just set to the DNA site type.

We also need to place particles onto the lattice to represent the transcription start site for each gene in the genome. First, the transcription start and end sites are assigned to individual 10 bp beads in the configuration. The genomic sequence positions were obtained from the NCBI GenBank entry for Syn3A: CP016816.2. While the transcription start site typically is found upstream of the coding sequence (CDS), we take the genomic sequence position of the CDS for each protein-coding gene as the start and end sites for transcription.

#### Morphology: RDME Lattice Site Types

The RDME simulation requires site types to be defined for every position in the cubic lattice to evaluate diffusion propensities and determine reaction localization. We first initialize a simulation with a simulation box of dimensions 64 64 128 in units of lattice sites whose edges lengths are 10 nm. The size of the z-dimension is twice the size of x and y because the simulation box will contain two cells divided along the z-axis after division. With the simulation box initialized, the cellular regions can be defined using the RegionBuilder function in jLM. RegionBuilder contains functions to build filled predefined shapes (e.g. spheres and cubes) as binary arrays with dimensions equal to the size of the simulation box. It also contains functions to manipulate the binary arrays by growing or reducing regions. Because the regions are first defined as binary arrays, they can be manipulated using logical operations. First, we define a cytoplasmic sphere with radius 20 lattice cubes (200 nm) centered at the midpoint of the simulation box (32,32,64). From this sphere, we use the dilate function of RegionBuilder to create a sphere one lattice site thicker using the se26 option. The dilate options are se6 if contiguity of the new layer is not important or se26 if contiguity needs to be maintained in the new layer of lattice sites. 6 corresponds to the number of faces for adjacent cubes and 26 includes all edge and vertex adjacencies. We then use a logical operation to exclude the cytoplasm from the new sphere by using the “&~” (and not) operation between the two binary arrays. This leaves a spherical shell that defined the cytoplasm peripheral to the membrane with region name outer cytoplasm. From the first dilated sphere, we then add one more layer of lattice sites using the same procedure and then exclude both the cytoplasm and outer cytoplasm to leave a spherical shell for the membrane. Everything outside of the membrane is defined as extracellular. Once all region shapes are constructed (including the ribosomes and DNA), they need to be combined into the RDME site lattice. Because there will be overlaps between regions (e.g. between cytoplasm and DNA), an order of priority is assigned to each site type. The priority order from lowest to highest priority is as follows: extracellular, cytoplasm, outer cytoplasm, ribosomes, DNA, ribo centers, and membrane. This order is assigned so that only the membrane is exposed to the extracellular environment and so that the DNA excludes ribosome diffusion into the DNA, but does not exclude the center of mass chemically active site from performing translation.

### Simulation Methods for Chemical Reactions

#### ODE

Ordinary differential equations (ODEs) model the time evolution of metabolite concentrations in a deterministic framework. This method was applied to processes such as glycolysis, nucleotide synthesis, lipid synthesis, and transporter activity. We solved the resulting stiff ODE system using the LSODA solver from the ODEPACK software suite, employing the backward differentiation formula (BDF) method with order varying between 1 and 5.^57,58^ The system of ODEs was constructed using the odeCELL package, which provides a simple API for specifying kinetic parameters, defining custom enzyme kinetics rate forms, and mapping reactions to differential equations.

#### CME

The chemical master equation (CME) captures the stochastic dynamics of well-stirred systems by modeling the probability distribution over discrete particle counts and simulating chemical reactions as probabilistic transitions between states. Unlike the deterministic ODE model, which reports only mean concentrations, the CME approach captures both distributions and fluctuations, which is especially important in systems with low copy numbers. This method was used to simulate transcription and tRNA charging. Stochastic simulations were performed using the Gillespie direct algorithm as implemented in version 2.5 of Lattice Microbes (LM). The CME system was constructed using the pyLM package, which provides Python bindings to the underlying LM codebase.

#### RDME

The reaction-diffusion master equation (RDME) extends the CME by incorporating spatial resolution, dividing the system into subvolumes where reactions occur and between which particles diffuse. Simulations of the RDME sample trajectories from the underlying probability distribution *P* (**x***, t*), where the state vector **x** encodes the copy numbers of all species in all subvolumes:

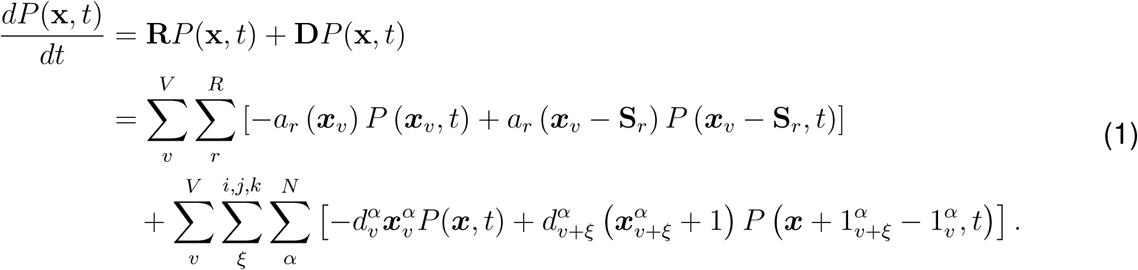

Here, the first term **R** corresponds to the CME description, which accounts for reactions occurring within each subvolume *v* according to the stoichiometric matrix **S** and reaction propensities *a_r_*(**x***_v_*) for each of the *R* reactions specified by *r*. The second term **D** is the diffusion operator, where *d^α^* is the diffusion propensity of species *α* in volume *v*, with *ξ* indexing the six Cartesian directions for the six neighboring volumes.

We used the RDME framework to simulate spatial and stochastic processes including RNAP diffusion and binding to gene start sites, translation, diffusion and binding of mRNA and degradosomes during mRNA degradation, membrane protein insertion, and replication initiation. Diffusion is described in further detail below. Importantly, RNAP and degradosomes were assigned distinct diffusion coefficients to reflect their differing mobilities from other proteins, and ribosomes diffuse within a special scheme that represents the ribosome on the lattice. Simulations were conducted using the multi-particle diffusion algorithm in Lattice Microbes v2.5.^59,60^ The spatial model and reaction network were constructed using the jLM package, which extends pyLM to support spatial modeling and interactive visualization.

### Hybrid Simulation Algorithm

The hybrid simulations are built on the platform of performing “hook” simulations in the single-GPU 32-bit RDME solver in LM IntMpdRdmeSolver. A summary of the hybrid simulation algorithm is shown in Figure 1. The hookSimulation function allows the user to interrupt the stochastic reaction-diffusion solver to perform user-defined functions through the jLM Python API. In a hookSimulation, the user can manipulate site types and the positions and numbers of particles in the RDME. Because the procedures exist inside a Python API, the user can execute any Python functions while the RDME solver is paused. The only restriction is that the hookSimulation function must return one of three recognized values by the LM solver: 0, 1, or 2. If the user returns a value of 0, nothing is changed in the RDME solver. If the user returns 1, the LM solver will read any manual changes made to the particle lattice and copy the updated particle lattice to the GPU. If the user returns 2, the solver also reads any changes to the site type lattice and writes the current state of the site types to the LM trajectory file.

The procedure for the 4DWCM hookSimulation function can be found in the deposited code for the model in Hook.py. The frequency of interruption to perform a hookSimulation is a userdefined variable at the start of the simulation in units of RDME time steps. Here, our hook frequency is 250 time steps, or 12.5 ms of biological time. This frequency was determined by the frequency required to update ribosome positions (discussed in Diffusion methods). Because we are communicating between spatially-localized and well-stirred reaction methods, we chose to track all particle counts dynamically in a single Python dictionary. The instantaneous particle counts and some state variables (e.g. volume and surface area) are dynamically updated throughout communication and are converted to and from concentrations only within the ODE model for metabolism. To communicate the different methods at their own frequencies, we simply implemented conditionals that follow timers for individual processes. The timer for communicating with BD chromosome dynamics is set to enter DNA communication every 4 s of biological time. Once the hookSimulation passes the DNA communication conditional or functions, it enters into the procedure to update the ribosome excluded volume. Ribosomes are updated after DNA because the ribosome excluded volumes depend on the excluded volume of the DNA. A timer is set for 1 s of biological time for the global CME and metabolic ODE simulations. The simulation first enters into the global CME procedure, reads the current cell state from the RDME, executes a 1 s simulation of well-stirred stochastic reactions for transcription and tRNA charging, and then the results are read from the CME trajectory into the particle counts and RDME lattice. The cell state is then communicated to the ODE model of metabolism where metabolites, transporters, and metabolic enzymes’ particle counts are first converted to concentrations using the instantaneous cell volume. The costs of genetic information are communicated to the metabolite pools. We then integrate 1 s of biological time using LSODA as described above. The results are communicated to the overall particle counts. Finally, also at an interval of one second, we write out the cell state. The particle counts dictionary and metabolic fluxes are recorded to a csv file. We save the particle and site lattice as binary npy files in the case that we need to restart the simulation from the current cell state. If the simulation is at an interval of one second, the hookSimulation function returns 2 to the LM solver to write the RDME state to the LM trajectory file. Otherwise, we return 1 so that the updated ribosome excluded volumes are copied back to the GPU.

#### Computational Expense of the 4DWCM

The average computational expense for each of the major components of the 4DWCM are plotted in Figure 1. The component requiring the most time is the RDME solver itself performing diffusion and spatially-localized stochastic reactions on the lattice. Second, is the time to update ribosome positions. In order to maintain excluded volume and have the ribosomes obey Stokes-Einstein diffusion, we update the projections of the excluded volume of every ribosome every 12.5 ms, or 80 times per biological second. The other notable contributor is the “DNA time”. While the time presented in the performance plot corresponds to the Growth and Division and Brownian Dynamics components of the flowchart, it does not include the time spent integrating the BD on the second GPU. The BD simulations are executed in parallel and take a similar amount of time per biological second compared to the total time spent in other processes. The time in the plot represents the time to perform communications, update morphology on the lattice, move particles to keep them in their respective regions (e.g. cytoplasm and membrane), and execute any serialized DNA processes. For example, the boundary in LM and LAMMPS must be synchronized, so we update the position of the membrane in LAMMPS serially instead of in parallel any time the morphology changes.

#### Simulating Chromosome Dynamics

In our minimal cell model, the chromosomal DNA is represented using a coarse-grained bead-spring polymer, implemented using the code btree chromo which calls LAMMPS as a library. Chromosome dynamics are simulated in hook intervals corresponding to 4 seconds of biological time. This approach builds upon the methodology introduced by Gilbert et al.^6^, enabling us to model the influence of SMC complexes and topoisomerases. In this section, we briefly review the core aspects of the original model and highlight the key modifications and extensions required to adapt it to the 4DWCM.

#### Polymer Representation and Brownian Dynamics

Syn3A’s chromosome is modeled as an elastic worm-like chain, with a coarse-graining of 10 base pairs per bead, with each bead having diameter *σ*_DNA_=3.4 nm.^27^ Bonds between DNA beads are finite extensible nonlinear elastic (FENE) using bond style fene in LAMMPS:

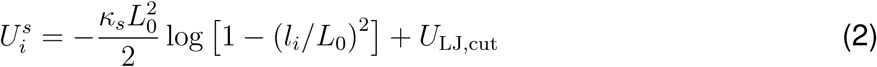

where *l_i_* is the distance between DNA beads *i* and 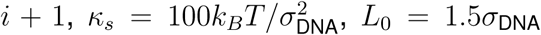, and *U*_LJ,cut_ is a Lennard Jones potential with *ɛ* = *k_B_T* and *σ* = *σ*_DNA_. The bending potential is implemented using angle style cosine:

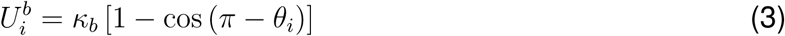

where *θ_i_* is the angle between DNA beads *i* and *i* + 1 and *κ_b_* = *l_p_k_B_T/σ*_DNA_ with persistence length *l_p_*=45 nm.^61^

Excluded volume effects are implemented via a “hard” potential which does not allow strand crossing, and “topo” potential which allows strand crossing. The hard Lennard-Jones potential is implemented with pair style hard,

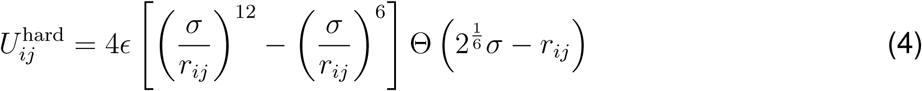

where *r_ij_* is the distance between beads *i* and *j*, *σ* = *σ*_DNA_, and the potential strength is *ɛ* = 5*k_B_T*. Every five looping cycles (i.e. every two seconds of biological time), we replace the hard potential with a “topoisomerase” potential which is implemented with pair style soft:

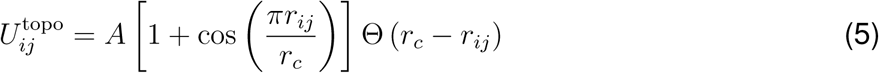

where *r_c_* = *σ*_DNA_ is the cutoff distance, and the potential strength is *A* = 0.1*k_B_T* or 1.0*k_B_T*. When simulating topoisomerase action, we perform the following sequence: 1) minimize with *A* = 0.1*k_B_T* and harmonic bonds, 2) Minimize with *A* = 0.1*k_B_T* and FENE bonds, 3) Run 5000 BD steps with *A* = 0.1*k_B_T* and FENE bonds, 4) Minimize with *A* = 1.0*k_B_T* and harmonic bonds, 5) Minimize with *A* = 1.0*k_B_T* and FENE bonds. Steps 1-3 allow strand crossings, and steps 4-5 ramp up the potential to avoid clashes when simulating with the hard potential.

Excluded volume between the DNA and the cell membrane is implemented with a hard potential, with the boundary particles representing cell membrane shapes being kept fixed during all minimizations and BD. For our model, we do not implement any attractive forces between the DNA and boundary particles, which could arise through translation-transcription coupling.

The system evolves under Brownian dynamics, modeled as Langevin dynamics in the large friction (no mass) limit. The equation of motion for each bead is given by 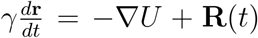 where *γ* is the translational damping constant, *U* represents the total interaction potential, and **R**(*t*) is a stochastic noise term satisfying the fluctuation-dissipation theorem, with an assumed temperature of 310 K. We use a timestep of 0.1 ns for integration. Time stepping was performed using a custom version of fix brownian that was modified to work with the Kokkos package in LAMMPS (see “GPU Acceleration” below). The translational damping constant for the DNA was calculated using the Stokes-Einstein equation *γ* = 6*πηr*_DNA_, with a dynamic viscosity *η* of 1.2 Pa s, which is derived from mRNA diffusion in *E. coli* as described in our previous 4DWCM.^1^

#### SMC Looping Model

We employ a simple model of SMC looping with one-sided extrusion and a 4 second residence time, using the methodology of Bonato and Micheletto,^62^ as implemented by Gilbert et al.^6^ Loop extrusion by an SMC dimer involves one domain anchoring to the chromosome while another domain acts as a hinge that extrudes DNA while the anchor remains anchored in place on the chromosome. At the start of each 4 second hook interval, anchor beads are selected randomly and uniformly across the chromosome, emulating SMC binding non-specifically with the DNA. Hinge beads are selected 6 beads away from the anchor in a random direction along the polymer. Anchor and hinge pairs are connected by harmonic bonds 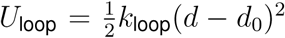 where *k_loop_* is the loop stiffness and *d*_0_ is the preferred loop size. For our simulations, *d*_0_ is set to 4*r*_DNA_ and *k*_loop_ is set to 0.33*k_B_T*. Anchor-hinge bonds are relaxed by sequence of two energy minimizations; the first with harmonic bonds between DNA beads, and the second with FENE bonds between DNA beads. Every 0.4 seconds (ten times every 4 second hook interval), the hinge beads are updated by the SMC step size, which we take to be Poisson distributed with a mean of 20 beads and truncated at a maximum of 30 beads. The step size of roughly 200 bp, and step frequency of 1 per 0.4 seconds, are based on values observed from in vitro experiments of yeast condensin, where the rate of loop extrusion by SMC complexes was observed to take place at rates up to 600 bp/s, with individual step sizes of the complex having a median around 200 bp.^23,24^ When choosing the new hinge bead, only beads whose distance from the anchor is less than 50 nm (roughly the size of a fully extended SMC complex) are considered. In btree chromo, SMC looping is performed by calling the function simulator run loops. For each 0.4 s biological time interval, we run 10000 BD steps, corresponding to 1 *µ*s of simulated time.

This study employed simplified SMC–SMC and SMC–replisome interaction dynamics. SMC proteins were allowed to extrude past one another, but Hi-C data from *B. subtilis* are best reproduced by simulations in which SMCs can block each other and, in some cases, facilitate each other’s unloading.^49^ Similarly, extrusion past replication forks was prohibited in our model, although there is evidence that occasional bypassing occurs.^50^ We also do not explicitly model interactions between SMC and RNA polymerase, although RNAP can act as a moving barrier that significantly impedes SMC extrusion at highly transcribed operons.^51^ Additionally, we assumed a fixed dwell time of 4 second and assume that all SMCs remain bound during that 4 second dwell time. The binding and unbinding kinetics of SMC proteins remain poorly characterized, but it is likely that the actual dwell time exceeds 4 seconds and that only a fraction of SMCs are bound at any given time. Finally, while SMC proteins extrude unidirectionally in our model, recent studies have shown that SMCs can switch direction, effectively acting as two-sided extruders,^52^ and simulations incorporating two-sided extrusion, rather than one-sided, better reproduce the juxtaposition of bacterial chromosomal arms.^63^

#### Partitioning of Daughter Chromosomes

Fluorescence imaging in this study shows that daughter chromosomes are spatially segregated into distinct cell volumes by the onset of FtsZ-ring constriction, indicating that partitioning occurs rapidly following replication. Since the precise mechanisms underlying chromosome partitioning remain unknown, we treat partitioning as a geometric constraint in our model by applying a repulsive force between daughter DNA strands to ensure segregation into distinct daughter volumes. We applied a repulsive partitioning force of magnitude 12 pN (corresponding to 5 *k_B_T/*1.7 nm), active for each of the 10000 × 0.1 ns BD timesteps during each loop cycle of 0.4 biological seconds. In btree chromo, this force can be toggled using the command switch fork partition repulsion:T. This force was applied only to already-replicated DNA and restricted to a spherical region of 2000 Å centered between the daughter cells. A 12 pN force is applied to every DNA bead in each daughter chromosome, directed along the unit vector connecting their centers of mass. This vector is computed at the beginning of each 0.4 s interval, and the forces are applied in opposite directions to drive segregation. It was deactivated (switch fork partition repulsion:F) once chromosomes were fully partitioned (i.e., both chromosomes had passed the mid-plane and entered their respective daughter volumes), which prevented the partitioned chromosomes from being pressed against opposite sides of the cell.

The 12 pN partitioning force used in our simulations exceeds thermal forces, which are typically considered to be below 10 pN.^64^ However, because we apply frequent energy minimizations and simulate only 1 *µ*s of Brownian dynamics for each 0.4 seconds of biological time, the force magnitude is decoupled from real-time dynamics. In our simulations, the partitioning force serves as a constraint to ensure reliable separation of the daughter chromosomes into their respective cellular volumes without inducing excessive compaction at the cell poles, and does not represent a quantitatively calibrated value based on the underlying physics or biological mechanisms.

#### Omission of Potential Biological Mechanisms Contributing to Partitioning

It is still not fully understood which mechanisms contribute to chromosome segregation and partitioning in various cellular contexts. Proposed drivers include SMC- and topoisomerase-mediated loop extrusion and disentanglement, and entropic forces.^26,65^ Although our simulations explicitly model SMC complexes and topoisomerases, they lack sufficient Brownian dynamics (BD) timesteps to capture entropic segregation, were it to occur. The “fork partition force” discussed above emulates daughter-daughter repulsion that could arise from entropic effects. To further complicate things, recent findings suggest that entropic forces may hinder, rather than promote, segregation in some contexts,^66^ underscoring the need for out-of-equilibrium dynamics.

BLAST searches of the Syn3A genome indicate that it lacks a ParAB*S* system, as it does not contain genes encoding either ParA or ParB, nor does it possess complete signature *parS* sites. The ParABS system, which is present in *Bacillus subtilis* and *Caulobacter crescentus*, promotes chromosome alignment and segregation by preferentially loading SMC at *parS* sites near the origin of replication.^48^ In E. coli, the SMC complex MukBEF loads preferentially on newly replicated DNA and is excluded from the Ter region by the MatP/matS system,^67^ and the Min system aids in chromosome partitioning through MinD mediated transient DNA-membrane tethering events,^47^ neither of which we were able to identify in Syn3A. We were also unable to identify genes encoding PopZ and TipN, which, in *C. crescentus*, function alongside the ParABS system to anchor the chromosome to the membrane and facilitate partitioning.^68^ In *Mycoplasma pneumoniae*, superresolution imaging suggests anchoring of the Ter-proximal region of the DNA to the attachment organelle.^69^ Not only does *Syn3A* lack orthologs of genes for chromosome-partitioning proteins, but cryo-electron tomograms of Syn3A also show no visible attachment organelle or analogous structures.^21^ In contrast, cryo-ET of *M. pneumoniae* shows a distinct membrane-associated attachment organelle.^40^ In the minimal cell, there is no evidence of preferential loading or unbinding for SMC, nor has there been direct observation of DNA-membrane anchoring. As part of our efforts to model partitioning mechanisms, we tested an origin-origin repulsive force inspired by DNA-membrane tethering systems found in other bacteria but this had negligible impact on partitioning and was not pursued further. We also did not model transcription-translation coupling, although it could plausibly assist in partitioning through functioning as DNA-membrane tethering. It has also been proposed that contributions from polysomes may aid in chromosome partitioning.^70^ As discussed earlier, polysomes were not included in our model.

#### Omission of the Twisting Potential

In contrast to Gilbert et al., we do not include a twisting potential in our representation of the DNA polymer. Although supercoiling plays a critical role in genetic information processing by regulating transcription and influences chromosome conformation through the transient formation of localized writhe in the form of plectonemes, given the coarse-grained resolution of our model (10 nm cubes), the effects of supercoiling are expected to have a small affect on the chromosome conformation when projected onto the lattice. Consequently, we do not explicitly model supercoiling generated by complexes such as SMC-ScpAB, DNA gyrase, RNA polymerase, or DNA polymerase. Furthermore, the 1 kb resolution 3C-Seq contact map that we have for Syn3A indicates that there is no persistent supercoiling, meaning that if plectonemes do form, they must be transient and rapidly resolved.^21^

The Kokkos package, described below, does not currently support the polytorsion potential used in the Gilbert et al. methodology, which used aspherical particles and quaternions to represent the DNA beads as rigid bodies. Additionally, energy minimizations in LAMMPS do not adjust quaternions, meaning torsional degrees of freedom remain fixed during minimization. While it is possible to reintroduce torsional angles using dihedral interactions with phantom beads, this approach was not pursued in the current study to maintain computational efficiency while focusing on large-scale chromosomal organization.

#### GPU Acceleration

Simulations were conducted using LAMMPS with the Kokkos package to leverage GPU acceleration. The Kokkos backend allowed efficient parallelization of Brownian dynamics and energy minimizations. For this study, custom Kokkos versions for fix brownian and fix addforce were written, based off of the existing Kokkos versions of fix langevin and fix setforce already in LAMMPS. For our system size of approximately 100K beads, using LAMMPS with Kokkos on the Delta A100 GPUs improved performance by about a factor of 10 compared to CPU-only execution.

#### Communicating Chromosomes Between LM and LAMMPS

The state of the DNA on the LM RDME lattice depends directly on the BD chromosome state, but also communicates cell state information to the LAMMPS simulations. In the direction of the chromosome model to the RDME, the coordinates of the 10 bp beads determine the lattice sites in the RDME that are assigned the “DNA” site type and determine where chemically active particles (e.g. gene start sites, the origin of replication particle) are located on the lattice. Reassigning the site types first involves reassigning all intracellular lattice sites that were previously DNA to cytoplasm. Subsequently, each lattice site containing any monomers from the new chromosome configuration are assigned as a DNA lattice site. Following the site type update, chemically relevant DNA particles are moved on the lattice. Based on their positions in the genome, corresponding particles in the chromosome configuration are mapped to individual 10 bp particles. Their current state (e.g. whether a gene has an RNAP bound or not) and position is checked on the RDME lattice and that particle is deleted. A particle in the same chemical state is then placed in the lattice site containing the 10 bp bead corresponding to that particle’s position in the genome. Communication with the membrane boundary is discussed in the morphology Methods.

We determine the communication time to the chromosome based on the loop extrusion rate of SMC. As mentioned above, we selected to communicate with the chromosome to update their positions every 10 loop extrusion steps, corresponding to 4 seconds of biological time. As input to btree chromo, we count the instantaneous number of SMC proteins and use that information to determine the total number of loops that are simulated. This number is calculated simply as half of the number of proteins from the gene *smc*/0415, rounded down, and is written to the input file of btree chromo as the integer loop count in the function simulator run loops. At the communication time, once the information is communicated from the BD chromosome model to the 4DWCM, the next 4 s of biological time are set to run as a background process on a second GPU using the Python subprocess.Popen functionality. Running as a background process on a second GPU means that the rest of the 4DWCM can continue with the chromosome dynamics being calculated in parallel, eliminating a wait time of 2-5 minutes per 4 seconds of biological time. As discussed below, updating division morphologies are instead performed serially.

#### DNA Replication

We previously simulated DNA replication in the stochastic well-stirred kinetics where replication forks work read through entire genes all-at-once.^1^ Because the BD chromosome structure is at a resolution of 10 bp, we also increased the resolution of replication kinetics to 10 bp increments. The replication forks independently traverse the opposite arms of the mother chromosome while replicating the DNA, with beads for the left and right daughter chromosomes appearing in locations centered about the location of the mother’s corresponding bead, consistent with the so-called “train-track” model.^26^ As a simplification, we assume that both the leading and lagging strands are replicated simultaneously. The replication topology (the position of the replication forks at 10 bp resolution) is tracked explicitly in the BD chromosome model. Using this information, we calculate the rate of replication in front of each replication fork independently. We continue to use the replication rate of 100 bp/s, and the chromosome state is updated in the RDME at an interval of 4 seconds, meaning the maximum amount of replication per fork per communication is 400 bp. We determine the rate of replication given the instantaneous pools of dNTPs using the template-driven, enzyme catalyzed rate form for DNA replication^71^ (excluding the binding step)

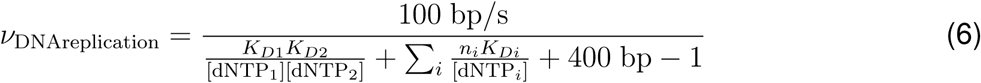

where the input sequence is the 400 bp in front of the respective replication fork. Using the instantaneous dNTP pools from the metabolism as input, the polymerization rate gives us an instantaneous rate of replication that determines the number of base pairs (divided by 10 to determine the number of Brownian dynamics beads) that are replicated at the communication time. In btree chromo we update the replication state using the transform function whose input variables are the number of DNA beads to add at each replication fork using the train-track model described above. Because the chromosome state in the RDME is read directly from the state in the BD simulation, we do not need to manipulate the replication state on the RDME lattice.

### Chemical Reaction Models

#### Metabolism

The set of metabolic reactions remains unchanged from the well-stirred model simulated in Thornburg et al., 2022.^1^ The systems biology markup language (SBML) reaction model is read into the model using cobrapy to set the reaction set and stoichiometries. We store kinetic parameters in an Excel file (see Table S2) that has forward and reverse catalytic rate constants, a Michaelis-Menten constant for each substrate and product, and an enzyme annotation for each metabolic reaction. The complete set of kinetic parameters for the metabolic reactions along with the free energy calculations for central, nucleotide, and lipid metabolism are provided in the the Supplemental Information. The changes in parameters from the previous model are described below.

In glycolysis, we increased the forward rate of the fructose bisphosphate aldolase reaction (FBA) from 21.0 s^−1^ to 64.5 s^−1^, both of which are reported values in BRENDA.^72^ Two parameters were changed in the phosphorelay reactions for glucose uptake. First, we increased the rate of PtsI binding to phosphoenolpyruvate (PEP) (reaction GLCpts0) from 6,000 mM^−1^s^−1^ to 10,000 mM^−1^s^−1^. Both of these values come from kinetic measurements performed on the same protein in *E. coli*.^73^ We also increased the rate of PtsG taking up and phosphorylating a glucose molecule (reaction GLCpts4) from 0.88 mM^−1^s^−1^ to 1.29 mM^−1^s^−1^. The new rate constant was calculated from the glucose uptake rate measured in *M. pneumoniae* with the corresponding PtsG proteomics count and external glucose concentration.^74^

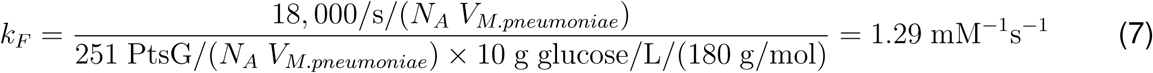

The combination of increased glucose uptake and increased rate of the FBA reaction resulted in all cells surviving for the entire cell cycle. We chose to simulate the set of parameters in which more cells are healthy to demonstrate that there is a solution to the set of parameters in which the entire population of simulated cells is healthy. In the absence of a measured death rate for Syn3A, it is unclear which parameter set better mimics the real cell fates of Syn3A.

The other metabolic parameters changed were the uptake rates of (deoxy)nucleosides. In our previous well-stirred model, we observed excess build-up of (d)NTPs. We do not account for inhibitory mechanisms in the metabolic kinetics, which could result in sufficient uptake rate reductions such that the cell does not build up excess pools of nucleotides. In the absence of inhibition, we instead reduce the following uptake rates: adenosine, guanosine, uridine, deoxyadenosine, and deoxyguanosine uptake rates were reduced from 2.0, 1.0, 2.0, 1.5, and 1.0 s^−1^ to 1.5, 0.5, 1.5, 1.0, and 0.5 s^−1^, respectively.^1^

#### Replication Initiation

Replication initiation as previously simulated in the well-stirred stochastic kinetics representation. With spatially-localized DNA particles, we now move this set of reaction to the RDME representation. The following reactions are based strongly off of our previous reaction models.^1,36^ In transcription, RNAP particles diffuse to chemically-active promoter particles on the RDME lattice whose positions follow their respective monomer in the BD chromosome. Similarly, we will treat the origin (*oriC*) as a single RDME particle whose position follows the origin monomer in the BD chromosome. Although the replication initiation conglomeration will be larger than the size of a single 10 nm lattice site, the replication initiation protein will only look for a single position relative to the origin at any one time. Therefore we assume that because we are not concerned with the excluded volume of the origin that we can represent the origin as a single particle that undergoes many state changes as proteins bind. The protein responsible for initiating bacterial DNA replication is DnaA. DnaA has two DNA-binding domains: domain IV to double-stranded DNA and domain III to single-stranded DNA. Replication initiation begins by a single DnaA protein binding to a 9/9 consensus sequence (TTATCCACA) with a high affinity (HA) near *oriC*. In our model this is modeled as a single binding reaction:

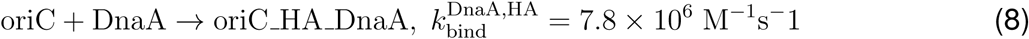

We assume that this binding is irreversible. Following the first DnaA, there are two nearby 7/9 consensus sequences within 5 bp of the HA site. We assume that DnaA binding to these with its dsDNA binding domain with a low affinity (LA). We assume that the order of binding to these two sites does not matter and that the DnaA will bind irreversibly to them one after the other.

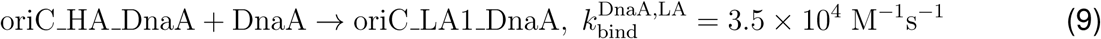

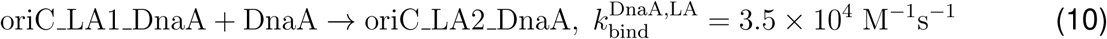

Following the first three DnaA bound to the dsDNA sites near oriC, we assume that a small amount of torque is applied to the DNA opening up a small amount of ssDNA in the known AT-rich region directly neighboring the dsDNA binding sites.^36^ DnaA is known to bind cooperatively and also to ssDNA, so we simulate the remainder of replication initiation as a reversible polymerization reaction where DnaA bind to the end of the filament along the DNA one DnaA at a time. These reactions follow the scheme

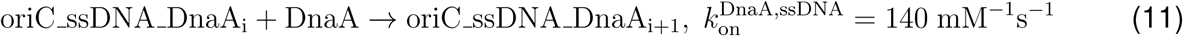

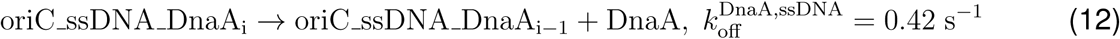

where *i* is the number of DnaA bound to the ssDNA (excludes the 3 bound to dsDNA). Based on the length of the AT-rich region, we assume that the filament can consist of up to 30 DnaA. However, while DnaA is the initiation protein, its responsibility is to open the origin so that the replication machinery can be loaded. The final reaction of replication initiation is the loading of the replisome. Based on the size of a complete replisome as the most conservative estimate, we assume that the machinery can be loaded once there are 20 DnaA in the filament so that a large enough pocket of ssDNA has been formed. A minimum of 20 corresponds to roughly 20 nm of linear DNA. Based on a molecular reconstruction of a bacterial replisome (PDB: 5IKN),^75^ we assume 20 nm of linear DNA is sufficient to load the first replisome. We simulate the replisome loading reaction using a representative protein, DNA helicase DnaB. We assume that if the DnaB finds an unwound origin, it will be immediately recruited to assist in replication, so we simulate the binding as a set of fast reactions

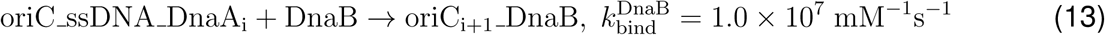

where *i* is the number of DnaA in the filament ranging from 20 to 30. We keep the number of DnaA in the species name so that we can track the number of DnaA bound to *oriC* at the time of replication initiation. We assume that two replisomes are required, so *oriC* undergoes this reaction twice. However, once a single DnaB binds, we assume that the state of the origin particle is irreversible and also that no more DnaA need to bind.

In the phase of replication initiation when DnaA are forming a filament along ssDNA, we first used the parameters from our previous well-stirred model: 100 mM^−1^s^−1^ for binding and 0.55 s^−1^ for the on and off rate, respectively.^1,36^ In early testing, we found that these rates were not sufficient to initiate replication in the 4D model. The spatial localization adds the restriction that DnaA must diffuse to the *oriC* particle, affecting the overall rate of polymerization of DnaA. The new rates favor stronger binding of DnaA to ssDNA: 140 mM^−1^s^−1^ for binding and 0.42 s^−1^. Both sets come from the same single molecule FRET study.^41^ The original parameters represent the average values measured in the experiments and the new parameter set represents a set with a bias toward stronger DnaA binding when the ssDNA is preceded by dsDNA containing the strong and intermediate signatures.

#### Transcription of mRNA and tRNA, Translation and mRNA Degradation

The transcription, translation, and degradation reactions were modeled as binding events between macromolecular polymers (DNA, mRNA) with their respective machinery (RNA Polymerase, Ribosome, and degradosome), followed by (de)polymerization reactions. Binding in the RDME occurs when the molecules diffuse into the same lattice cube and subsequently interact. These reaction kinetics are implemented similarly to our previous model^1^, which provides a detailed account of the underlying methods. Here, we briefly summarize the methods and focus on key changes introduced in the present work.

The binding rates of polymers and corresponding macromolecular machinery were estimated based on the frequency of the binding events reported for *E. coli*; each case will be discussed individually below. The stochastic binding reactions follow a simple bimolecular form (A+B to C). In the RDME (Eq. 1), the propensity *a_r_*, which quantifies the probability per unit time that reaction *r* will occur, is calculated for a second order binding reaction as 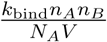, where *k*_bind_ is macroscopic binding rate reported in units of M^−1^s^−1^, *n_A_* and *n_B_* are the absolute counts of molecules A and B respectively, *N_A_* is Avogadro’s constant, and *V* is the volume. Over long times, the average number of reaction events per unit time equals the time-averaged propensity, and we approximated the propensity by the observed frequency of binding events. Rearranging, we obtain the formula for the macroscopic binding rate *k*_bind_:

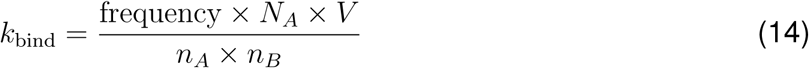

The Hofmeyr rate form^71^ is applied to catalyzed, template-directed polymerization of RNAs and proteins as described in our previous work.^76^ For RNA synthesis (i.e. transcription elongation), the enzyme and template pair is RNAP and DNA, and for protein synthesis (i.e. translation elongation), the pair is ribosome and mRNA. The formula for the Hofmeyr rate—excluding the step where the enzyme binds to the template polymer—takes the form:

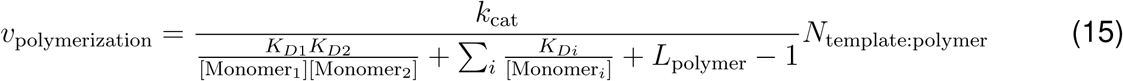

Here, *k*_cat_ is the intrinsic elongation speed of enzyme on the template in units of length (bp/nt/aa) per second. *K_D,i_* are the dissociation constants of monomers of type *i* in the polymer (e.g. ATP, CTP, UTP, and GTP for transcription). The concentrations of monomers are dynamic over the cell cycle, and shortages—which occur if consumption overtakes synthesis—can slow or in some cases even halt chain elongation. For transcription, the monomers are NTPs with *K_D,_*_NTP_ of 0.1 mM, and for translation, monomers are aa:tRNAs with *K_D,_*_aa:tRNA_ of 1 *µ*M.^36^

First, we describe the parameterization for transcription reactions, starting with the transcription of tRNA-coding genes. Due to the lack of measurements for RNAP binding with tRNA-coding genes, we instead take the value measured for rRNA-coding genes. We use the values of 10 initiations of transcription of rRNA genes per minute and 11,400 RNAP molecules in a single *E. coli* as measured by Bremer et al.^77^ The volume of a single *E. coli* cell is taken as 1fL, so the binding rate between tRNA genes with RNAP 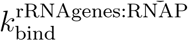 was calculated as

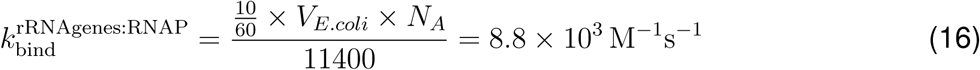

Thus, the binding between tRNA-coding genes with RNAP occurs via

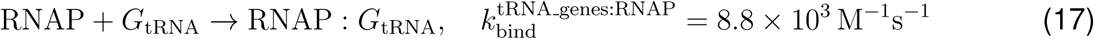

The elongation speed of RNAP on tRNA-coding genes is taken as 85 nt per second, which is the value measured for RNAP on rRNA-coding genes in *E. coli* ^77^. The elongation occurs via

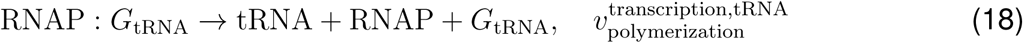

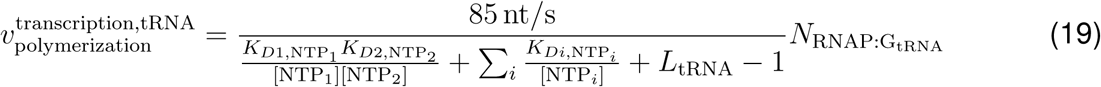

A promoter strength 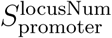 was introduced as a prefactor at the transcription level to distinguish the expression strengths of the 455 different mRNAs. It was incorporated into both the binding and the elongation reactions to account for variations in both RNAP binding affinity and transcription elongation speed. Promoter strengths were scaled according to the absolute protein counts,^76^ using the formula

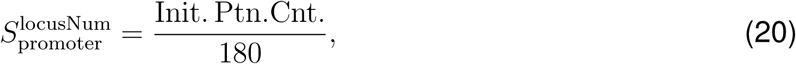

where 180 is the average count of the 455 mRNA-coding proteins, and the initial protein count as described in the section *Initial Conditions*. Ribosomal proteins use proxy promoters strengths corresponding to 500 proteins if their experimental proteomics value was less than 500. If their experimental proteomics count is greater than 500, that value is used instead.

The average binding rate of RNAP with mRNA-coding genes 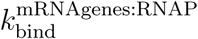 was scaled as

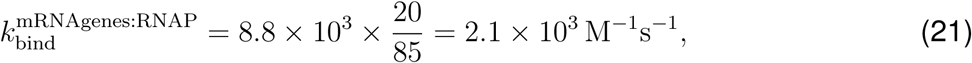

since the average elongation speed of RNAP on protein encoding genes is taken as 20 nt/s.^78^

The binding rate of RNAP with each unique gene 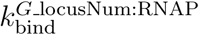 is

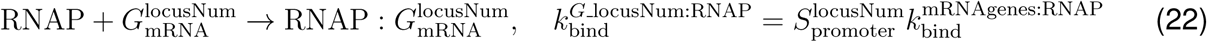

The elongation speed of RNAP on mRNA-coding genes is assumed to scale linearly with the promoter strength 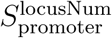 as 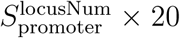 nt/s, where 85 nt/s is assigned as the upper limit.

Then, the polymerization of mRNA proceeds as

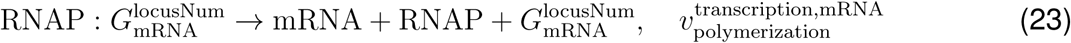

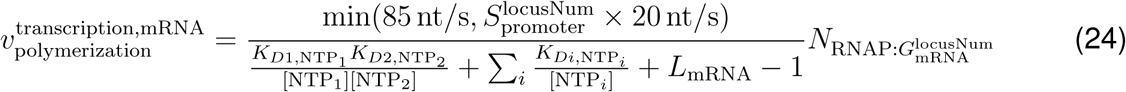

Translation was conducted in a similar fashion as transcription. Reported values for the mean time for translation initiation per mRNA vary widely, with a minimum of 1 second and median of 15 seconds.^79^ In our model, we assume a translation initiation frequency of 40 events per minute. The number of ribosomes in slow-growing *E. coli* has been experimentally measured as 6800.^80^ Then the binding rate between mRNA and ribosome 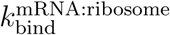 was calculated as

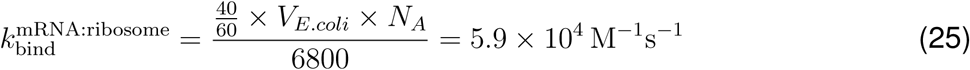

The translation speed was set to 12 aa/s.^81^ The synthesis of proteins happens via the binding and elongation reactions

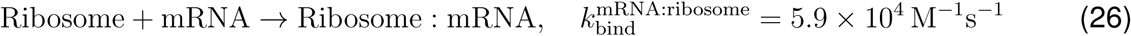

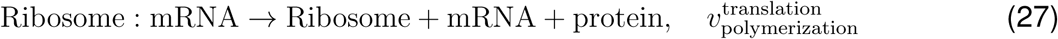

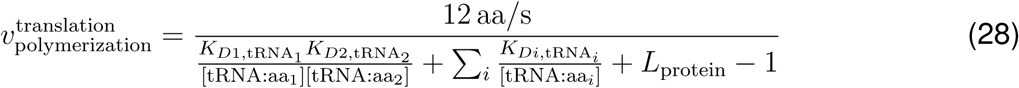

Finally, we describe the parameterization of mRNA degradation. The degradosome is localized in the peripheral membrane region via its scaffold RNase Y, which anchors the complex through its N-terminal transmembrane helix embedded in the membrane^82^. Therefore mRNA must diffuse to the periphery in order to bind with the degradosome. The binding rate between mRNA and degradosome 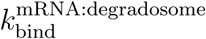 is estimated based on the observed value of 11 cleavage events of RNase E per minute per RNase E^83^, and 7800 mRNAs^84^ measured in *E. coli*.

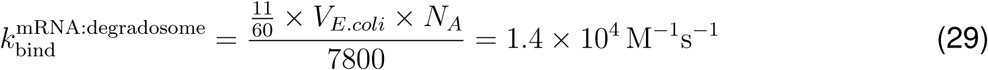

The degradation rate of an mRNA polymer chain was considered constant over the length of the chain^76^, with the speed taken to be 88 nt / s^85^. The product of mRNA degradation is a collection of nucleotide monophosphates. The respective reactions are

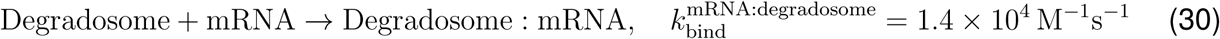

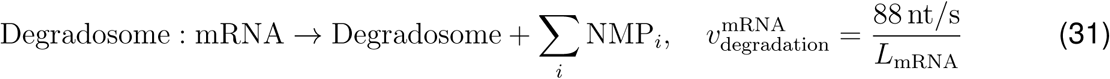

ATP energizes bond formation in tRNAs and mRNAs and breaks down mRNAs at the expense of one ATP molecule per monomer. Two GTP molecules are required in protein chain elongation for each amino acid, one for the delivery of aa: tRNA to the A site, and another for the shift of the ribosome to the next codon.

#### Simultaneous Transcription of rRNA Genes by Multiple RNAP

While we do not handle polycistronic RNA/transcription of multiple genes in operonal structures, we found that we will never generate sufficient rRNA to double the number of ribosomes in a cell cycle unless we allow multiple RNAP to read the genes coding for rRNA simultaneously. We handle this in a manner similar to the origin particle for DNA replication initiation where we have single particles that undergo state changes that track the number of molecules associated with the particle. In this case, the gene particles associated with the rRNA genes undergo the state changes based on the number of RNAP transcribing the gene. We assume a RNAP spacing of 400 nt, so the 5S genes can only have one RNAP at a time, but the 16S and 23S genes can have up to 4 and 7 RNAP at once, respectively.

The kinetics are implemented as follows. On the RDME lattice, the genes start in free states where no RNAP are bound. Once an RNAP diffuses and binds to the gene particle through the reactions described above, the gene is switched to a state indicating that 1 RNAP is bound to the gene and that the start of the gene is occupied. In the global CME kinetics, the transcription reactions are not for the entire gene like all other genes. Instead, we transcribe the gene in increments of 400 nt. Once the RNAP has transcribed the first 400 nt of the gene, we switch the state of the gene particle on the RDME lattice to indicate that the start of the gene is now unoccupied even though there is an RNAP still actively transcribing downstream on the gene. This particle can now react with a second RNAP in the RDME to initiate a second simultaneous transcription event. In the global CME, the transcription of this RNAP is also simulated in increments of 400 nt. However, we restrict the transcription of the second RNAP so that it cannot pass the transcription state of the first RNAP. The resolution of this traffic is limited to 400 nt because of the resolution of the individual transcription reactions. So once the first RNAP advances to its next 400 nt increment, the second RNAP can advance past the start site, once again freeing the start site of the gene. Because of this implementation, we also allow for gaps in transcription because the timescale between transcription initiation reactions is stochastic. RNAP progress in 400 nt increments until the end of the gene where the last transcription elongation reaction in determined by the remaining length of the gene sequence. As mentioned above, we allow RNAP to initiate on rRNA genes until all 400 nt increments are fully occupied.

#### RNAP and Degradosome Assembly

The assembly of RNAP and degradosomes were handled in a simple fashion. For the RNAP, we assumed the complete complex to be two alpha subunits *rpoA*/0645 and the two different beta subunits *rpoB*/0804 and *rpoC*/0803 (beta’). Our assembly assembly pathway assumes the two alpha units bind first, then the beta, and finally the beta’. For these reactions we assumed a fast binding rate of 10^7^ M^−1^s^−1^. The degradosome is a complex that consists of several proteins, some of which are metabolic proteins whose roles in the degradosome are unknown.^86^ For this reason, we assume the assembly of a minimal number of proteins to constitute a “complete” degradosome: the scaffold protein and endoribonuclease RNase Y *rny* /0359 and an exoribonuclease RNase J1 *rnjA*/0600. The binding of these two proteins uses the same fast binding rate as the RNAP assembly reactions.

#### Ribosome Assembly

Assembly of the ribosome is more complicated due to the tens of proteins and 3 rRNA species involved in the complex. We base our parameters on the assembly of the 30S small subunit (SSU) in *E. coli*. The kinetic parameters come from pulse-chase experiments.^87,88^ In the kinetic model for ribosome biogenesis, we determined a hierarchical association of rproteins to the RNAs forming the small subunit. An assembly tree of intermediates at two different temperatures was constructed based on fluxes in the reactions of individual proteins/ribosomal RNA interactons.^87,89^ The hierarchical pathways were consistent with the in vitro assembly map assigning primary, secondary, and tertiary rproteins.^90,91^ Because of the tens of proteins involved in the complex, there are combinatorially many ways to assemble the SSU. Primary rproteins bind directly to the 16S rRNA and secondary and tertiary proteins bind once other proteins have bound. The number of assembly intermediates in the tree can be trimmed to enhance computational performance. An assembly tree of only 145 intermediates was obtained by choosing reactions that were observed to have flux above a predetermined threshold. We further reduced this branched tree to a linear pathway consisting of only 19 intermediates in which only one rprotein was added at a time. We found that the assembly time was not significantly affected in the volume of Syn3A with this reaction set, with assembly times still being less than 1 minute after a 16S rRNA is transcribed. We additionally found that the variations in diffusion coefficient for each intermediate throughout assembly did not significantly affect assembly time, so we have assigned uniform diffusion coefficients for all intermediates to the diffusion coefficients of their respective rRNA backbone. We also also assume a linear reaction scheme for the assembly of the 50S large subunit (LSU). This reaction scheme follows the binding order reported by in *E. coli*.^88,92^ Because the rate constants for each binding reaction have not been as precisely quantified as the SSU assembly reactions, we assume two approximate binding rates: one weak binding rate (10^4^ M^−1^s^−1^) and one strong binding rate (10^6^ M^−1^s^−1^). These values were estimated from the orders of magnitude for weak and strong binding reactions in the SSU assembly. Whether a ribosomal protein (or the 5S rRNA) are a weak or strong binder was assigned using the weak/strong binding strength assignment in the LSU assembly maps for *E. coli*.^88,92^ Syn3A’s genome codes for two rRNA operons. While the isolated rRNA particles are fully differentiable, we do not differentiate assembled ribosomes or SSU/LSU intermediates based on rRNA isoform.

#### tRNA Charging

The reactions to charge tRNA are simulated in the global CME representations. We use the global CME because tRNA and aminoacyl tRNA synthetases (aaRS) are particles that are low in count whose states are dynamically changed by small molecules that are high in concentration and fast diffusers: ATP and amino acids. We assume a reaction scheme that begins with the aaRS binding with an ATP.

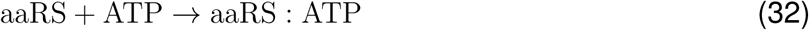

We then assume the aaRS:ATP complex binds with the corresponding amino acid.

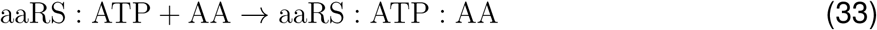

We do not account for the possibility of the aaRS binding to the incorrect amino acid. The aaRS:ATP:AA then binds to an isoform of its respective tRNA.

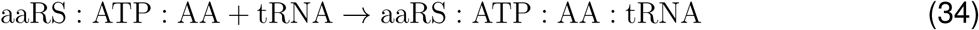

Again, we do not allow for the aaRS to bind to the incorrect type of tRNA. While there are 20 tRNA types, Syn3A has redundant isoforms for some with a total of 29 tRNA-coding genes.^36^ Finally, we simulate a conversion reaction where the aaRS releases the charged tRNA, an AMP, and a di-phosphate.

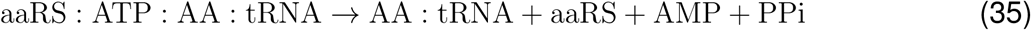

To communicate the costs of translation to the charged tRNA pools, we treat the cost counters like particles in the well-stirred global CME. Instead of simply subtracting the cost counters for translation from the charged tRNA pools, the cost particles undergo fast reactions with their respective charged tRNA to convert them into a “paid” state.

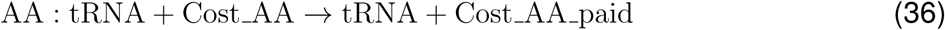

Because we are communicating the cumulative cost per biological second, the total cost of translation almost always exceeds the total amount of charged tRNA because we only have roughly 200 tRNA per isoform. If we were to use the method of subtracting the cost from the available pools, we would almost always artificially have remaining costs that are not paid. By allowing the cost counters to be paid one amino acid at a time through a fictitious reaction, the translational AA:tRNA costs are dynamically paid off every biological second.

There are two special cases in tRNA charging: the formylmethionine and glutamine tRNAs. For glutamine, the charging mechanism involves mischarging the tRNA with a glutamate and then replacing it with a glutamine.^18^ Here, we instead simulate a lumped reaction that mimics the above charging mechanisms, but uses the glutamyl-tRNA amidotransferase protein in place of the aaRS. Accounting for the mischarging reactions explicitly would require a custom algorithm to track the mass balance of the aaRS and tRNA states. While this is possible, it is not a targeted interest in this model, so we chose to summarize its effects with a lumped reaction. We do not explicitly simulate the reaction to convert the charged MET:tRNA MET into FMET:tRNA MET. Instead, we account for the formylated methionine tRNA by accounting for its cost in two places: the MET:tRNA MET and the metabolite 10fthfglu3. We add the FMET:tRNA cost to the MET:tRNA cost to account for the use of the methionine in the global CME. To account for the formylation, we treat the substrate for the methionine-tRNA formyltransferase reaction (10fthfglu3) as the pool to pay for the cost and communicate with it in the same way we communicate with other metabolic pools.

### Diffusion

#### Proteins and RNA

The diffusion rate of all cytoplasmic proteins is set to a conservative estimate of 1 *µ*m^2^/s given the dense crowding of Syn3A’s cytoplasm.^21^ To account for excluded volume of DNA, the diffusion into DNA lattice cites for cytoplasmic proteins is reduced to 0.5 *µ*m^2^/s because we estimate that the DNA occupies roughly half a lattice site on average.^21^ For ribosome excluded volume, we assume that proteins can exit the center lattice site of ribosome projections, but are forbidden from re-entering the center sites. The remainder of the cross projection is a slight overestimate of the excluded volume, so we assume a reduced diffusion rate of 0.5 *µ*m^2^/s for proteins into and out of the cross. We assume a diffusion coefficient of 0.1 *µ*m^2^/s for transmembrane proteins. Cytoplasmic proteins are forbidden to enter the membrane lattice sites and transmembrane proteins are forbidden to leave membrane lattice sites. Peripheral membrane particles obey the same diffusion rates as cytoplasmic proteins. However, their locomotion is restricted so that once they diffuse into a peripheral membrane cytoplasmic lattice site, they stay in the peripheral membrane lattice sites (they do not re-enter the cytoplasm and cannot enter the membrane).

The diffusion coefficients of all RNA are calculated estimates based on hydrodynamic radius.^93^ We used a modified version of the Stokes-Einstein relation where the diffusion coefficient can be approximated as

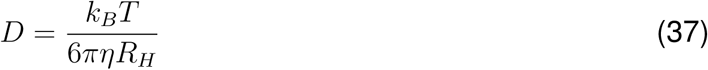

in which *η* is the viscosity of the environment and the hydrodynamic radius can be approximated as

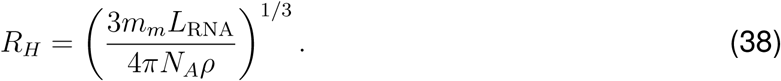

The viscosity was previously estimated to be 1.2 PA·s from the diffusion coefficient of an mRNA in *E. coli*.^1,94^ We assume a temperature *T* of 310 K, an average molar mass *m_m_* of 337 g mol^−1^ per nucleotide, and a density *ρ* of 1.8 g cm^−1^.^1^ With these values we calculate a length-dependent (*L*_RNA_) diffusion coefficient for every RNA. RNA particles are able to diffuse into any lattice site in the cytoplasm including DNA and ribosomes, but are forbidden from entering membrane lattice sites.

#### RNAP and Degradosomes

Once protein complexes assemble, they are free to diffuse within their respective simulation space (e.g. degradosomes in the peripheral membrane and RNAP in the cytoplasm). Degradosomes and RNAP are assigned diffusion coefficients of 0.031 *µ*m^2^/s and 0.22 *µ*m^2^/s respectively and are assumed to fit into a single 10 nm lattice cube. Diffusion of degradosomes is restricted so that they cannot leave the peripheral membrane layer of the cytoplasm.

#### Ribosomes

Ribosomes are roughly 20 nm in diameter, so they are considerably larger than a single 10 nm lattice cube and cannot be treated with normal diffusion in the RDME using 10 nm lattice cubes. To accurately represent a ribosome on the lattice, we utilize a center of mass particle around which we construct a 3D cross shape as shown in Figure S1. This projection of the ribosome is used to exclude the volume of the ribosome, influencing protein diffusion around ribosomes. The center of mass particle is also utilized as the reactive particle to initiate translation, so mRNA are allowed to diffuse into ribosome sites. Proteins are totally excluded from the center lattice site of the ribosome cross. The 7 lattice sites making up the ribosome lattice site projection overestimate the total volume of the ribosome, so we allow proteins to enter the exterior lattice sites of the cross at a reduced rate (~50%) of their cytoplasmic diffusion rate.

To allow ribosomes to diffuse on the lattice, we need to dynamically update the position of their projection on the lattice to correctly exclude their volume. The center of mass particle of ribosomes is assigned a diffusion coefficient of 0.001 *µ*m2/s and is only allowed to diffuse with the ribosome’s lattice sites. We assigned this diffusion coefficient based on the calculated diffusion coefficient for ribosomes from Brownian Dynamics simulations by Benjamin Gilbert where ribosomes were simulated as hard spheres with radii of 10 nm in the presence of a single chromosome within a cell of radius 200 nm.^6^ Gilbert et al. calculated a diffusion coefficient range of 0.01 0.03 *µ*m^2^/s for the ribosomes, however the simulations did not include the exact crowding that ribosomes would experience in Syn3A and instead used the estimated viscosity of *E. coli* cytoplasm from^1^, and Syn3A cytoplasm appears significantly more dense in cryo-electron tomograms^21^. As a conservative estimate, as Earnest et al. calculated when treating the dense nucleoid region of *E. coli* ^87,89^, we reduced the diffusion rate of the ribosomes by an order of magnitude within the dense cytoplasm of Syn3A to a value of 0.001 *µ*m2/s. With the diffusion coefficient and knowledge that the center of mass can diffuse only one lattice site (10 nm) without updating the excluded volume lattice sites, we can use Stokes-Einstein diffusion to determine the frequency at which the excluded volume sites need to be updated to maintain realistic diffusion. The minimum frequency can be calculated as

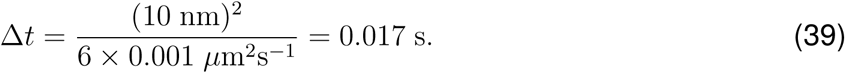

As a conservative estimate that also serves as a consistent frequency for all communication procedures, we chose to update the ribosome lattice sites every 12.5 ms or 250 RDME time steps.

Because each ribosome is independent in the RDME representation, they will diffuse apart even if they are translating the same mRNA. Adding polysomes then necessitates treating the diffusion of ribosomes in another computational method and communicating their positions to the RDME lattice. We attempted to add ribosomes as diffusing obstacles for the chromosomes in the Brownian dynamics LAMMPS simulations, but they caused frequent clashes with SMC loops. Unfortunately, even if the ribosomes were not causing clashing issues with SMC loops, the Brownian dynamics simulations still do not simulate enough time steps to reach the biological times to recover realistic diffusion. Therefore, addition of polysomes presents a significant challenge and will require integration of another method into the hybrid simulations.

### Cell Growth and Division in 3D

#### Calculating Cell Surface Area Dynamically from Lipids and Membrane Proteins

The surface area and volume in the simulations are directly tied to the chemistry of synthesizing membrane components. As lipids are synthesized and membrane proteins are translocated, we iteratively update the total surface area of the system. Each second of biological time, we count the number of lipids and membrane proteins, assign each molecule a surface area contribution per particle, account for the lipid bilayer, and then sum all the surface area contributions to get the total surface area of the membrane. We use the same assumptions as the previous well-stirred model for the individual surface area contributions in units of square nanometers per particle: membrane proteins - 28, cardiolipin - 0.4, sphingomyelin (sm) - 0.45, phosphatidylcholine (pc) - 0.55, phosphatidylglycerol (pg) - 0.6, phosphatidic acid (pa) - 0.5, Gal-DAG (galfur12dgr) - 0.6, 1,2-diacylglycerol (12dgr) - 0.5, and CDP-diacylglycerol (cdpdag) - 0.5.^1^ To account for the lipid bilayer difference, we estimate that for a spherical membrane 200 nm in radius, 51.3% of the lipids are in the outer monolayer. We assume that this fraction remains roughly constant throughout the cell cycle. The total surface area as well as the lipid and membrane protein contributions for the 50 simulated cells are shown in Figure S3.

To calculate the cell volume, we first make an assumption because we do not include the kinetics for FtsZ polymerization and neck constriction. We assume that the model grows spherically from its initial size until it approximately doubles in volume. Cells of roughly 200 nm and 250 nm (corresponding to an initial volume and a doubled volume) were previously observed in cryo-electron tomography, so it is a reasonable assumption that a cell might experience both these volumes in its cell cycle.^21^ In the simulation, we calculate the radius for a sphere with the surface area calculated using the surface area contributions discussed above and use this radius to calculate the cell volume during spherical growth. Once the volume doubles, we assume that it remains constant throughout the remainder of the cell cycle.

#### Updating Morphology During Spherical Growth

Once every 4 seconds of biological time, we update the total surface area and volume of the cell by calculating the total surface area as described above. During the period of the cell cycle when the cell grows to twice its volume, we assume that the cell grows spherically. Because we impose a cubic lattice on 3D space, our growth steps during this period are discrete even though we treat the surface area and volume as continuous quantities. From the instantaneous cell volume, we calculate the radius for a sphere with an equivalent volume. We update the morphology on the RDME lattice in increments of 10 nm because of the 10 nm lattice spacing. Therefore, we simulate membrane shape for cells with radii: 200, 210, 220, 230, 240, and 250 nm. In increments of 4 s of biological time, we check if the cell radius has passed a 10 nm threshold (following the frequency of updating the chromosome state to ensure the membrane boundaries are synchronized between methods). If a threshold has been crossed, the site types are updated on the RDME lattice to reflect the new cell size. To impose a new morphology on the lattice, we need to reconstruct the projection of each region onto the RDME lattice. The procedure for generating the regions are the same as generating the initial conditions. Once the new shapes are generated, we clear the site type lattice and reassign each lattice site to their updated site type. Ribosomes and DNA are left in place and we assume their dynamics will result in their exploration of the new space. Updating the positions of other particles during morphological changes is described below.

#### Generating Membrane Shapes During Division

We tested two methods of generating membrane shapes during division. The method that more closely reflected the morphologies observed in fluorescent imaging assumes that the cell divides symmetrically as two overlapping spheres (domes). We determine the radii and center-to-center distance for the domes geometrically using only the surface area and volume of the system. A 2D projection schematic is shown in Figure S3A. We can determine the volume of one of the domes by integrating discs. We assume that the center of the dome is the origin and integrate these discs for a dome of radius *R* laterally from *R* to *h* where *h* is the distance from the center of the dome to the division plane, or half the center-to-center distance between the domes.

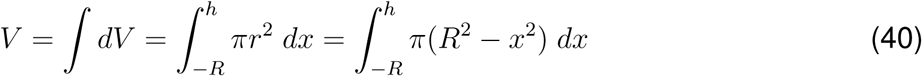

Evaluating the integral for volume gives us an equation relating the volume to the dome radius and distance to division plane.

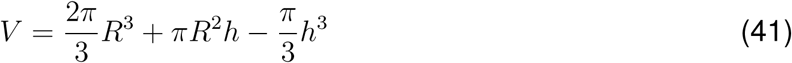

To connect the surface area to the dome radius and center-to-center distance, we use the surface of rotation method to rotate a 1-dimensional semicircular line about the axis of cell division. Our linear equation takes the form

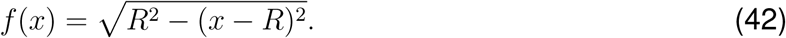

When we reduce the integral for surface of rotation, the integrand reduces to depend only on the dome radius.

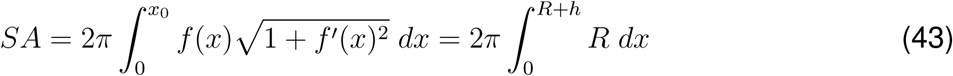

Evaluating this integral gives us an equation that relates the surface area to the dome radii and distance to division plane.

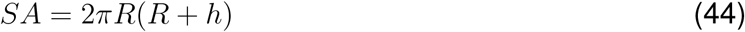

We do not solve this system of equations analytically. Once the cell volume has doubled in the simulation, we numerically solve for *R* and *h* using Equations 44 and 41 at a frequency of 4 seconds of biological time (following the frequency of updating the DNA configuration). When we solve for the parameters to determine cell shape, we double Equations 44 and 41 and set them equal to the instantaneous cell surface area and volume. The equations were solved for a single dome, but the cell consists of two mirrored domes making up the total surface area and volume. We could have instead halved the total surface area and volume of the system when solving to obtain the same results. The numerical solutions are obtained using the optimize module of the scipy Python library. After setting up the system of equations, we then use the least squares function to obtain the set of solutions. Because the equations are cubic, there will be multiple set of solutions and we need to choose the set that reflects the physical reality of the membrane shape. We know that the solutions that best mimics the cell shape are positive solutions where *R* is between its initial radius and the radius at doubled cell volume and *h* is between zero and the cell radius. We provide initial guesses of 0 to 300 nm to the least squares solver for *h* and *R*, respectively, to ensure we choose the realistic solution.

The values of *h* and *R* are then used to build two cytoplasmic spheres of radius *R* whose centers of mass are separated by distance *h*. The time-dependent values for *R* and *h* from the 50 simulated cells are shown in Figure S3C and D. In the solutions for *R*, we see that the cells increase in radius after they reach twice the initial surface area, indicating that the solutions do not perfectly preserve volume once the cell has doubled and divided. The division plane is defined as the midplane along the z axis of the simulation (z=64 lattice sites). We then define the cytoplasm of the dividing cell as the union between the two individual cytoplasmic spheres. The membrane is then constructed by building a contiguous layer of lattice sites at the surface of the cytoplasm using the dilate function implemented in jLM.

#### Updating Particle Positions During Morphological Changes

As the morphology is updated on the RDME lattice, the particles need to be moved to remain inside their respective regions in the cell. As the cell grows and divides, particles that are in the membrane must stay in the membrane and particles that are cytoplasmic need to stay inside the cell. There are many solutions to the problem, but because almost all morphological changes are minimal (radial changes on the order of a single lattice site), we chose a simple method that moves particles to the nearest lattice site of the type that they were in before the change.

For each site type in the simulation, excluding ribosome site types, we check each lattice site X that had site type Y before the morphological change. If lattice site X is still type Y following the morphological change, nothing is done. Otherwise, we use a K-Dimensional Tree (KDTree, scipy.spatial.KDTree function) as a binary mapping of the positions of all lattice sites of type Y following the morphological change. We then query the KDTree for the nearest lattice site to X that is type Y after the morphological change. Each particle from site X is placed into the new lattice site until all particles are placed or the site is full. If the site becomes filled, we once again query the KDTree for the next nearest lattice site of type Y and repeat the process until all particles from site X have been redistributed to the nearest sites of type Y. The algorithm for moving ribosomes is almost identical, but because they have their own lattice sites, we move them to the nearest cytoplasmic lattice site.

#### Communicating the Membrane Boundary to the Chromosomes

In communicating the RDME cell state to the chromosome configuration, the current shape of the cell is communicated to the program by determining the positions of the boundary particles in the btree chromo simulation. Before the onset of division, the membrane boundary is constructed using the spherical brdy function in btree chromo. The input variable is the instantaneous cell radius during spherical growth in units of Angstroms. The function creates a sphere of boundary particles who have repulsive interactions with DNA monomers. The boundary particles are packed on the surface such that there are no gaps for DNA monomers to escape through.

The process during division is more complicated, as instantaneous jumps in morphology can result in DNA monomers escaping the cell. The morphology during division is generated using the function overlapping spheres bdry whose input variables are the values of *R* and *h* discussed above (in units of Angstroms). This function generates a boundary for two symmetric spherical domes of radius *R* and separated by a distance *h*. The procedure generates a boundary similar to the spherical brdy function where boundary particles are packed tightly enough that DNA monomers cannot pass through. At time points when the morphology is updated, a custom procedure was required to ensure the DNA stays inside the cell. Instead of executing a LAMMPS simulation in parallel, a division procedure is executed serially because the RDME and BD membranes need to be synchronized. In this custom procedure, the initial condition is the input variables for *R* and *h* for the previous morphology. To guarantee that the DNA stays inside the cell, the boundary particles cannot be moved by more than the radius of a DNA bead. With this limitation, we compare the differences between the last *R* and *h* and the values for the new morphology. If either difference is greater than the radius of a DNA bead, the BD simulation is instructed to enter a progressive set of minimizations. We calculate the number of steps in increments of DNA bead sizes between the last and new values of *R* and *h*. The greater number of steps determines the number of minimizations. The procedure then iteratively creates membrane boundaries by incrementally changing *R* and *h* from the previous values to the new values and minimizes the energy of the chromosomes in each iteration. Because of the repulsive interactions between the DNA and membrane boundary particles, and because the membrane boundary particles are moved in increments less than the DNA bead size, the minimizations push the DNA monomers to the inside of the cell as the membrane constricts during division.

At the final step of division when the membrane fuses at the division plane separating the cytoplasms of the two daughter cells, an uncommon issue can arise where some DNA monomers can still be in the division plane. In this case, we defined a rescue procedure that clears all DNA from the division plane. To pull the DNA out of the division plane, we separate the two daughter chromosomes into independent BD simulations. The chromosomes are mapped to a daughter cell based on the positions of their centers of mass relative to the centers of the daughter cells. Once chromosomes are assigned to daughter cells, the pairwise distance between every DNA monomer in the chromosome and the center of the corresponding daughter cell is calculated. If the maximum distance is greater than the radius of the daughter cell, an iterative LAMMPS simulation is initiated. Again, boundary particles can be moved only in increments less than or equal to the radius of the DNA monomers. We initiate a simulation of the isolated chromosome in a spherical boundary whose radius is equal to the maximum calculated distance from before. The system is then minimized in progressively smaller spherical boundaries in radius increments equal to the radius of a DNA monomer until the membrane’s radius is equal to the radius of the daughter cell. These minimizations ensure all DNA monomers for the chromosome are out of the division plane and inside the cytoplasm of the respective daughter cell. Once both chromosomes undergo this procedure, their coordinates are recombined into a single LAMMPS simulation and the model continues with its normal procedures.

#### Generating Membrane Shapes with FreeDTS

In preliminary versions of the 4DWCM, we generated membrane shapes using the FreeDTS software to model the morphology of a growing cell by considering changes in three macroscopic characteristics: (i) an increase in volume, (ii) an increase in membrane surface area, and (iii) changes in the area difference between the two monolayers of the membrane. FreeDTS captures realistic membrane mechanics, as it uses the Helfrich Hamiltonian, which has been shown to accurately represent membrane behavior at large scales.^33^ However, under the parameters we tested, the resulting morphologies were more elongated—resembling teardrop shapes—rather than the nearly spherical geometries observed in fluorescence imaging. An example of such a teardrop morphology is shown in Figure S3. While this approach is more appealing, as it uses an accurate physical model of the membrane, we currently lack the necessary information and appropriate models for the interacting proteins that form the Z-ring, as well as a kinetic model of FtsZ polymerization and neck constriction. We expect that in the future, once these components are incorporated, the double-spherical shapes observed in experimental settings will be reproduced using a similar physics-based model. Therefore, we opted for the geometric approach as described above. Below, we briefly describe the original methods that were used to generate dividing membrane shapes.

FreeDTS has several potentials that can be used to influence membrane shape, and in our testing we used a Hamiltonian of the form

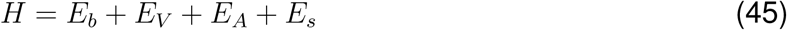

where *E_b_* is the bending energy of the membrane, *E_V_* is a potential dictating the total volume of a closed surface, *E_A_* is a potential dictating the total surface area of the system, and *E_s_* is a coupling potential to maintain global curvature of the membrane. In FreeDTS, the bending potential takes the form of a discretized version of the Helfrich Hamiltonian^95–97^ that is a function of the surface curvature up to the second order.

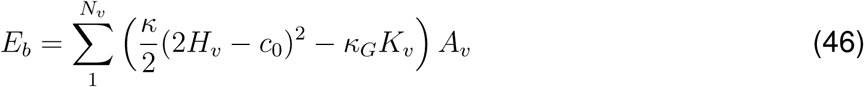

Here, the energy function is discretized to the energy evaluated at each vertex *v* in the surface. The mean curvature 2*H_v_* = *c*_1*,v*_ + *c*_2*,v*_, Gaussian curvature *K_v_* = *c*_1*,v*_*c*_2*,v*_, and area contribution *A_v_* are evaluated for each vertex, where *c*_1*,v*_ and *c*_2*,v*_ are the principal curvatures at the vertex. *κ* is the bending rigidity of the membrane, *κ_G_* is the Gaussian modulus, and *c*_0_ is the spontaneous curvature. Since the membrane surface did not undergo any topological changes, we safely ignored the second term, based on the Gauss-Bonnet theorem, as the total Gaussian curvature remains constant under pure deformations.^98^ We used a standard value for *κ* that is frequently used for simulating triangulated surfaces representing lipid bilayers: 30 k*_B_*T.^33,98^ The spontaneous curvature *c*_0_ describes how the membrane prefers to be curved. Because we assumed homogeneous membrane properties, this includes effects like the mismatch in area between sides of the lipid bilayer and deformation by membrane proteins. For example, a perfectly flat lipid bilayer has no area mismatch between leaflets, corresponding to a *c*_0_ of 0. In testing, we used the global curvature coupling potential (*E_s_*) instead, setting the coupling constant to 120 and curvature constant to 0.3, which were fixed throughout all of our simulations. This value was chosen because previous theoretical studies have shown that a curvature constant of 0.3 corresponds to dumbbell-like membrane shapes similar to the ones observed in imaging of Syn3A.^99^

While the parameters of the bending energy remain constant throughout our simulations, the variables come in the form of the surface area and volume of the cell as lipids and membrane proteins are incorporated into the membrane. FreeDTS has potentials to vary both the total surface area and volume of the membrane, however we will only be using the volume potential. Because we assume that the volume remains constant at twice its initial volume once the volume has doubled and continue to grow the surface area, it may seem intuitive that we should therefore utilize the area potential of FreeDTS, which takes the form

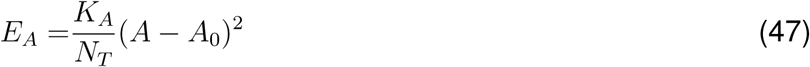

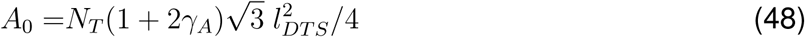

where *K_A_* is the compressibility, *N_T_* is the number of triangles in the surface, *l_DT S_* is the length units within a FreeDTS simulation, and *γ_A_* ranges from 0 to 1 to control the total surface area. In practice, topology is maintained in the simulation because the vertices have a volume exclusion between each other, and the edge lengths connecting them are constrained so that a vertex can never pass through a triangle (the edge length of a triangles can never exceed twice the diameter of a vertex). The constraints on the edge lengths are important to maintain constant topology, constraining the total surface and resulting in the range for *γ_A_*. The equation for *A*_0_ comes from the area of an equilateral triangle: 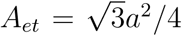 where *a* is the edge length of the triangle. In the range of values for *A*_0_, *γ_A_* = 1 corresponds to the case when the edge length of the triangle is twice the diameter of the vertex particles and *γ_A_* = 0 corresponds to the case when the edge length is equal to the diameter of the vertex particles (minimum because the vertex particles must still exclude each other). These constraints on the total surface area mean that it is difficult to fully explore the range of total surfaces areas that the cell experiences throughout growth and division without changing the number of vertices representing the membrane by adding more vertices to add more area. The dynamic addition of vertices has not yet been implemented into FreeDTS, so instead we constrained the total volume of the cell using the potential

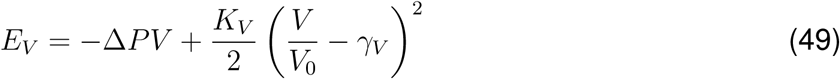

where Δ*P* is the pressure differential between the inside and outside of the membrane, *K_V_* is the compressibility, *V*_0_ is the volume of a sphere with the equivalent total surface area as the membrane, and *γ_V_* ranges from 0 to 1. The range for *γ_V_* represents the reduced volume of the system relative to the volume of a sphere with equivalent surface area. With this potential, it is recommended to only use the pressure term or the compressibility term, and not use both simultaneously. The pressure term should be used when you have knowledge of the pressure differential and not a target volume for the system. Similarly, the compressibility term should only be used if there is knowledge of a desired target volume for the system. This is because the compressibility term typically dominates the potential if it is used.^99^ We assumed the compressibility energy to be 10^5^ k*_B_*T, a value calculated for lipid vesicles^99^, and we consequently set Δ P equal to 0.

To generate division morphologies for Syn3A, we used FreeDTS to equilibrate triangulated vesicle shapes across a range of reduced volumes (*γ_V_* = 1.00 to 0.7). Starting with *γ_V_* = 1.00, we equilibrated the surface corresponding to a spherical membrane. This equilibrated shape served as the starting configuration for the next equilibration, using the same FreeDTS parameters but with *γ_V_* = 0.99. Equilibrations were then sequentially performed in *γ_V_* increments of 0.01, each equilibration using the final membrane state from the previous *γ_V_* as the initial conditions. Because we did not generate shapes that visually matched the experimentally observed division morphologies, we will not describe the preliminary methods that were developed to communicate these membrane shapes to LM and LAMMPS.

### Computational Analysis

#### Calculation of mRNA half-lives

Because we record the state of the lattice only once per biological second, we create book-keeping particles to track if gene expression events have occurred. If we do not explicitly track every gene expression reaction, there is a chance that reactions will not be accounted for in cost calculations communicated to metabolism. Additionally, we are able to accurately calculate bulk- behavior values like mRNA half-lives because we explicitly track all gene expression reactions to 1 second resolution. The following reactions create a book-keeping particle that is mapped to the gene locus for the individual RNA or protein: transcription, translation, mRNA degradation. For each process, the book-keeping particle is created as a product of the reaction. For example, the translation reaction for gene *dnaA*/0001 takes the form

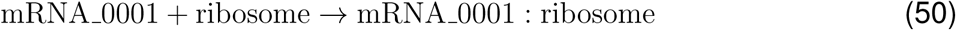

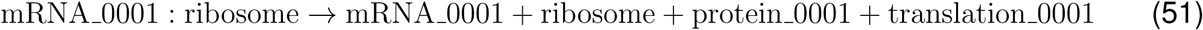

where translation 0001 is the book-keeping particle for translation of DnaA.

To calculate mRNA half-lives, we utilized the book-keeping particle for mRNA degradation. The overall rate of formation of a mRNA is

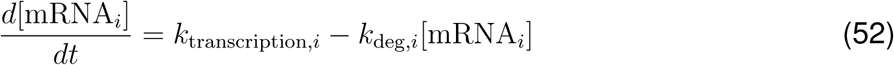

where the degradation rate *k*_deg*,i*_ is related to the half-life by

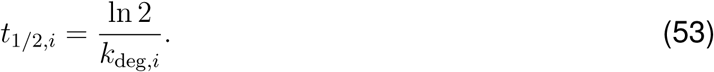

To determine half-life, we set the transcription rate to 0 and the overall formation rate equal to the total number of degradation events divided by the time elapsed in the simulation. These conditions reflect mRNA half-life experiments where transcription is halted and the counts of mRNA decrease as mRNA are degraded. Using the time-averaged concentration of mRNA, we can then calculate *k*_deg*,i*_ and half-life for each mRNA. The distribution presented in Figure 5 was calculated using the number of degradation events over 105 minutes.

### Computational Environments for Clusters and Supercomputing Resources

#### Docker Instructions

To build your environment using Docker version 20.10.11, build dea9396 (Docker Inc., U.S.), start by creating a Docker image based on the Nvidia CUDA 11.6.2 development container for Ubuntu 20.04 (nvcr.io/nvidia/cuda:11.6.2-devel-ubuntu20.04). Use the docker build command with your Dockerfile. This command automatically installs all necessary dependencies, such as the latest version of Miniconda (version 24.11.1, Anaconda, U.S., Linux-x86 64), along with make, wget, build-essential, libstdc++, libfmt, libssl, libssh, openssh-client, curl, libevent, zlib, manpages, and libsundials-dev (LLNL, U.S.), as well as Python packages including sbtab, pycvodes, pydantic, cobra, xlrd, and mpi4py. It also compiles additional software, including Lattice Microbes v2.5 (Zaida Luthey-Schulten Lab, UIUC, U.S.), odeCELL (Zaida Luthey-Schulten Lab, UIUC, U.S., https://github.com/Luthey-Schulten-Lab/Minimal_Cell/tree/main/odecell), and FreeDTS version 6.7.2023 (Niels Bohr Institute, University of Copenhagen, Denmark). To compile the software required for chromosome simulation, first install gen sc chain version 7.20.2023 (Zaida Luthey-Schulten Lab, UIUC, U.S., https://github.com/brg4/sc_chain_generation), followed by compiling GCC version 11.5.0 (GNU Project, international), CMake version 3.26.4 (Kitware Inc., U.S.), and OpenMPI version 4.1.5 (Open MPI Project, international). Next, compile LAMMPS version 19.Nov.2024 (Sandia National Laboratories, U.S.) and btree chromo (Zaida Luthey-Schulten Lab, UIUC, U.S., https://github.com/brg4/btree_chromo). Finally, run the Docker image interactively with the default conda environment activated using the docker run command.

#### Apptainer Instructions

To compile the environment using Apptainer version 1.3.5-1.el8 (Singularity/Apptainer Development Team, international), start by running apptainer build with your .def file, which uses the Nvidia CUDA 11.6.2 development image (nvcr.io/nvidia/cuda:11.6.2-devel-ubuntu20.04) as the base. Apptainer will copy the necessary source files into the container and then install core dependencies, including the latest version of Miniconda (version 22.11.1, Anaconda, U.S.), along with make, wget, build-essential, libstdc++, libfmt, libssl, libssh, openssh-client, curl, libevent, zlib, manpages, and libsundials-dev (LLNL, U.S.). It proceeds to build software, including Lattice Microbes v2.5 (Zaida Luthey-Schulten Lab, UIUC, U.S.), odeCELL (Zaida Luthey-Schulten Lab, UIUC, U.S., https://github.com/Luthey-Schulten-Lab/Minimal_Cell/tree/main/odecell), and FreeDTS version 6.7.2023 (Niels Bohr Institute, University of Copenhagen, Denmark). To compile the software required for chromosome simulation, first install gen sc chain version 7.20.2023 (Zan Luthey-Schulten Lab, UIUC, U.S., https://github.com/brg4/sc_chain_generation), followed by compiling GCC version 11.5.0 (GNU Project, international), CMake version 3.26.4 (Kitware Inc., U.S.), and OpenMPI version 4.1.5 (Open MPI Project, international). Next, compile LAMMPS version 19.Nov.2024 (Sandia National Laboratories, U.S.) and btree chromo (Zaida Luthey-Schulten Lab, UIUC, U.S., https://github.com/brg4/btree_chromo). Once completed, the Apptainer container will be fully prepared for execution.

## SUPPLEMENTAL FIGURES

**Figure S1:**
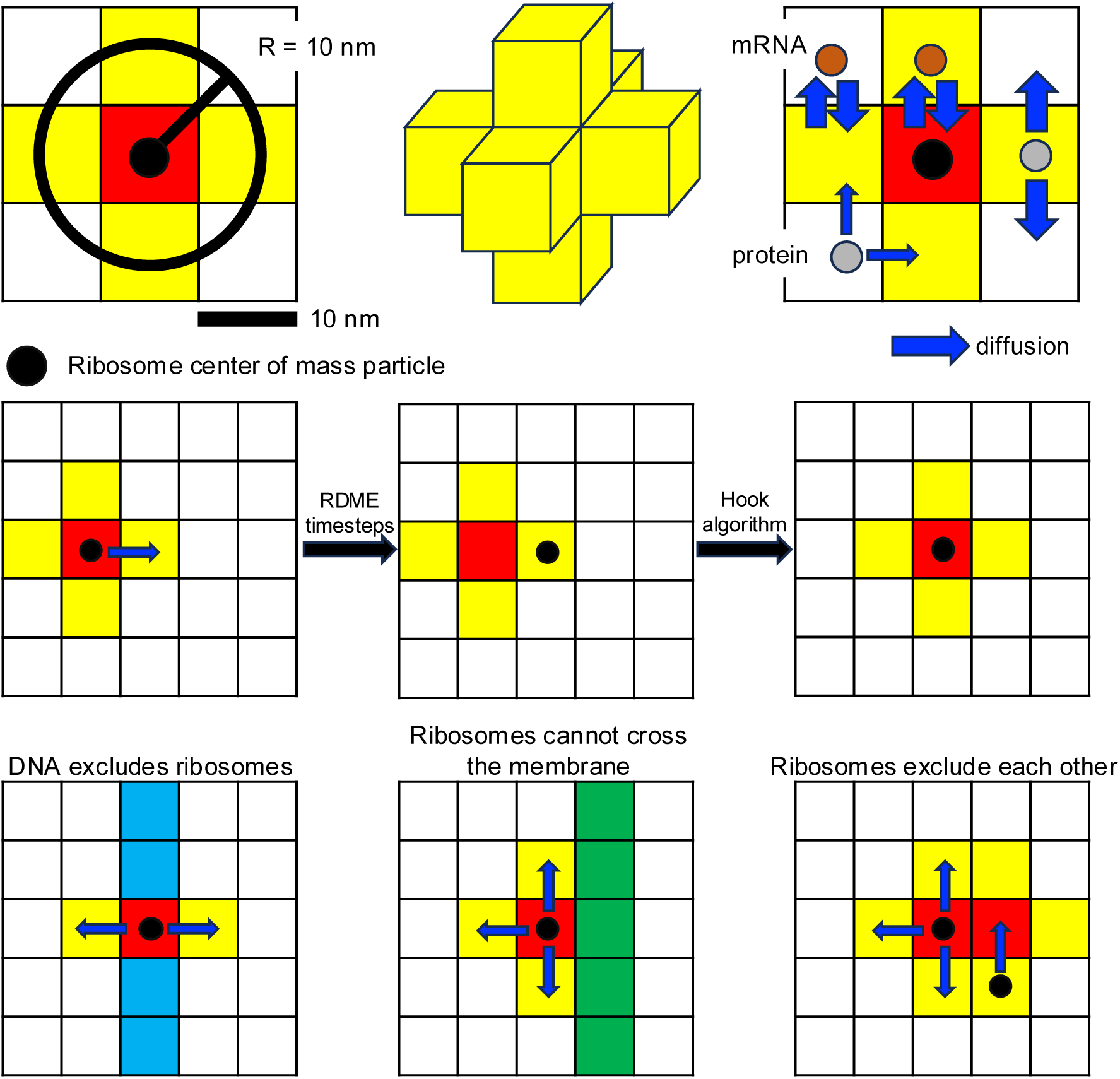
Projection of the ribosome onto the 10 nm lattice and its diffusion rules. The ribosome has a radius of 10 nm, resulting in the projections onto the lattice shown into the top left and middle. The top right shows that mRNA can diffuse into and out of ribosomes as their normal cytoplasmic rate so that mRNA can undergo translation initiation. Proteins can diffuse out of ribosomes at their cytoplasmic diffusion rate, but we reduce their diffusion rate into the ribosome because of the excluded volume of the ribosome. The center of mass particle of the ribosome starts at the center lattice site of the ribosome projection and is then allowed to diffuse within the cross. At the communication time, we update the position of the cross to reflect the new position of the center of mass particle.

**Figure S2:**
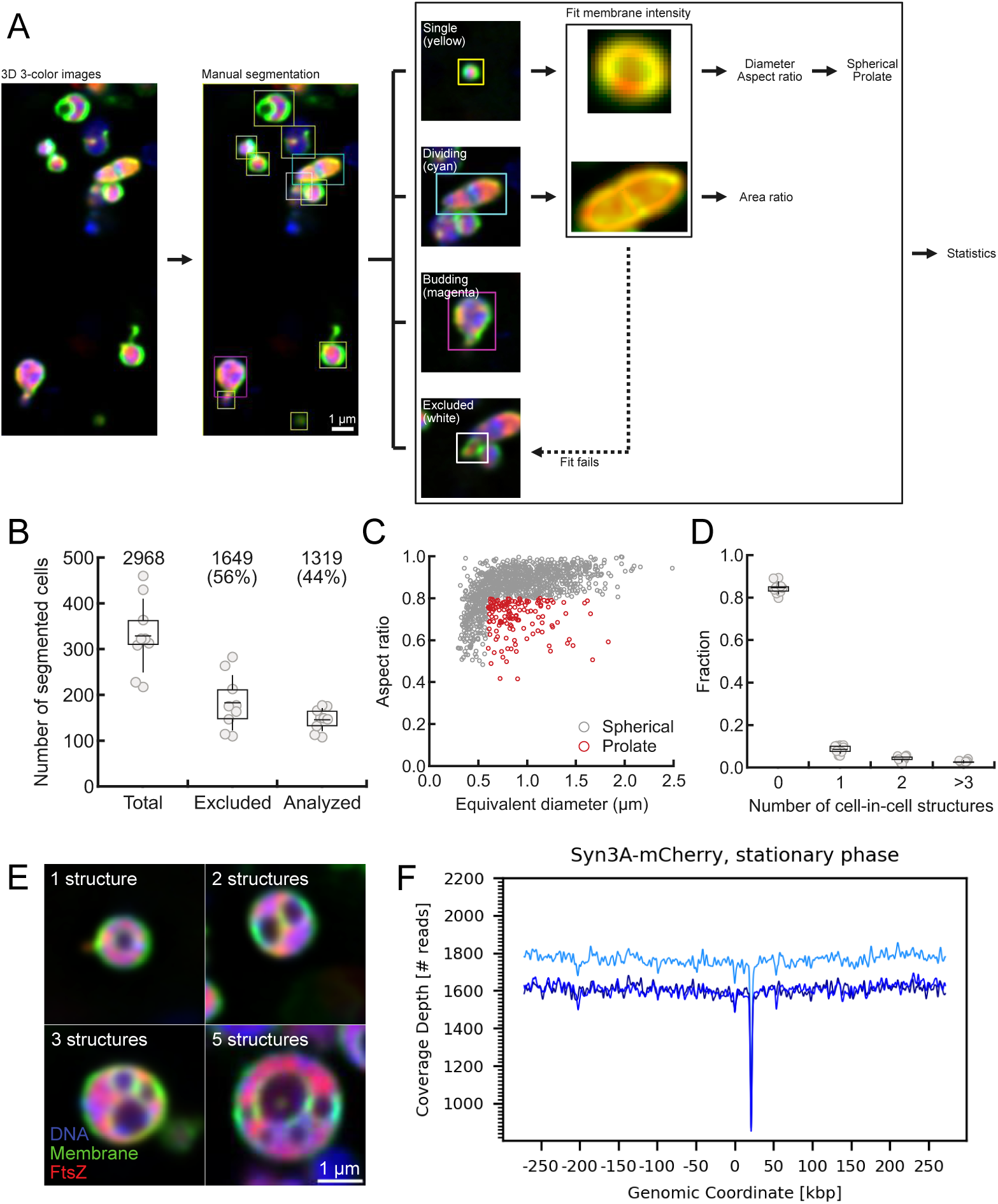
Imaging statistics and stationary-phase DNA sequencing of JCVI-syn3A/B. (A) Schematic representation of method to analyze morphologies in fluorescent imaging of JCVI-syn3B-FtsZ:mCherry. Color channels correspond to membrane (green), DNA (blue), and FtsZ (red). (B) Total number of segmented cells, the fraction that were excluded due to cell clusters, and the fraction that were analyzed and included in the morphology statistics. (C) Size and aspect ratio distribution of analyzed single cells. The average cell size was determined to be 0.89±0.37 *µ*m. The larger size is likely due to sample preparation method and lower spatial resolution in the imaging. Prolate cells were identified by quantifying the aspect ratio of their major and minor axes (see Methods). (D) Less than 20% of cells have “cell-in-cell” structures like the ones shown in (E). These have been observed previously in Syn3A and have been known to form in late exponential phase of growth.^19,32^ (F) DNA sequencing was performed in triplicate during the stationary growth phase. The lack of a slope in coverage corresponds to an ori:ter ratio of 1:1, indicating that the cells have stopped replicating. Each dot in a box plot represents a technical replicate (field of imaging) and covers the 25 to 75 percentiles. The means and standard deviations are represented as horizontal and vertical lines, respectively.

**Figure S3:**
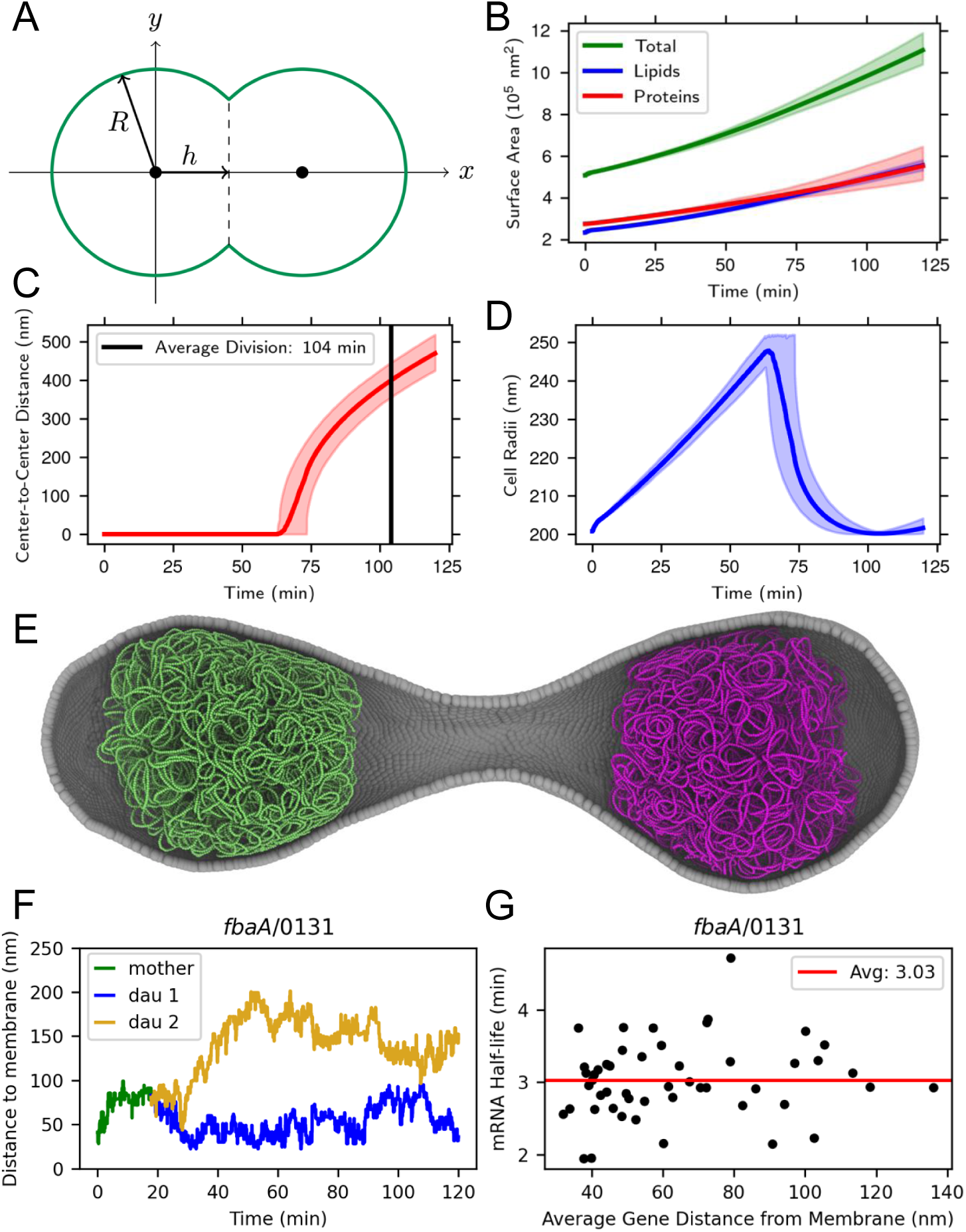
Spatial components of growth and division. (A) Schematic of the geometric division model. The dividing cell is treated as two overlapping spheres of radius *R* whose centers are separated by a center-to-center distance of 2 × *h*. (B) Total surface area and contributions from lipids and membrane proteins. (C) Center-to-center distance (2 × *h* from A) calculate from instantaneous cell surface area and volume. (D) Cell radii (*R* from A) throughout growth and division. The radius increases during spherical growth until division starts once the volume has doubled. In B, C, and D, lines show population average and shaded regions show full range among population. (E) Example of a membrane shape generated using FreeDTS. The long neck was not observed in experimental imaging. Grey particles represent vertices of the triangulated surface. Green and magenta particles represent two chromosomes fitted into the FreeDTS membrane structure. (F) Distance between the transcription start site particle on the RDME lattice for *fbaA* and its nearest membrane lattice site in a single simulated cell. At 20 minutes, the gene is replicated from the mother chromosome to the two daughter chromosomes. (G) Comparison of mRNA half-life to the average distance of the transcription start site particles for *fbaA* to the membrane. Each point represents a single simulated cell.

**Figure S4:**
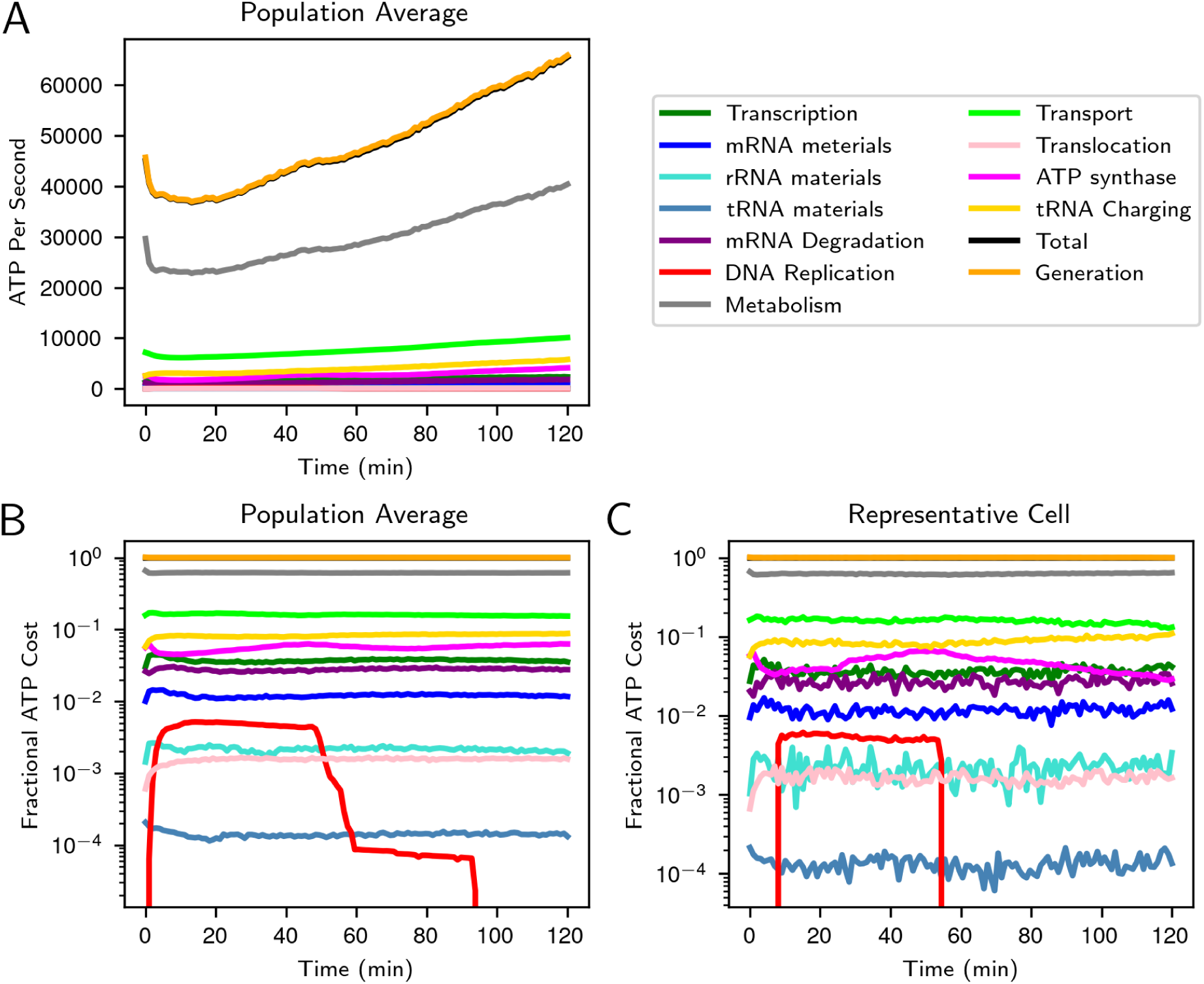
Cell-wide accounting of all ATP costs. Costs are averaged among the population (A,B) and for a representative cell (C). The total ATP generated is slightly higher than the total ATP cost to maintain the ATP pool as the cell grows and eventually divides. The cost of DNA replication is only present during replication, and the shoulders observed in the fractional cost plot (B) are a result the variations in the start and end of DNA replication among the population. The long plateau from 60 to 90 minutes is a result of the cell that initiated replication 46 minutes into its cell cycle, resulting in its replication to last until after 90 minutes. While ATP costs are steady on average, there are fluctuations in the costs in a single cell as observed in C.

## Notes

### Competing Interest Statement

The authors have declared no competing interest.

