## Supplemental Figures for "Bringing the Genetically Minimal Cell to Life on a Computer in 4D"

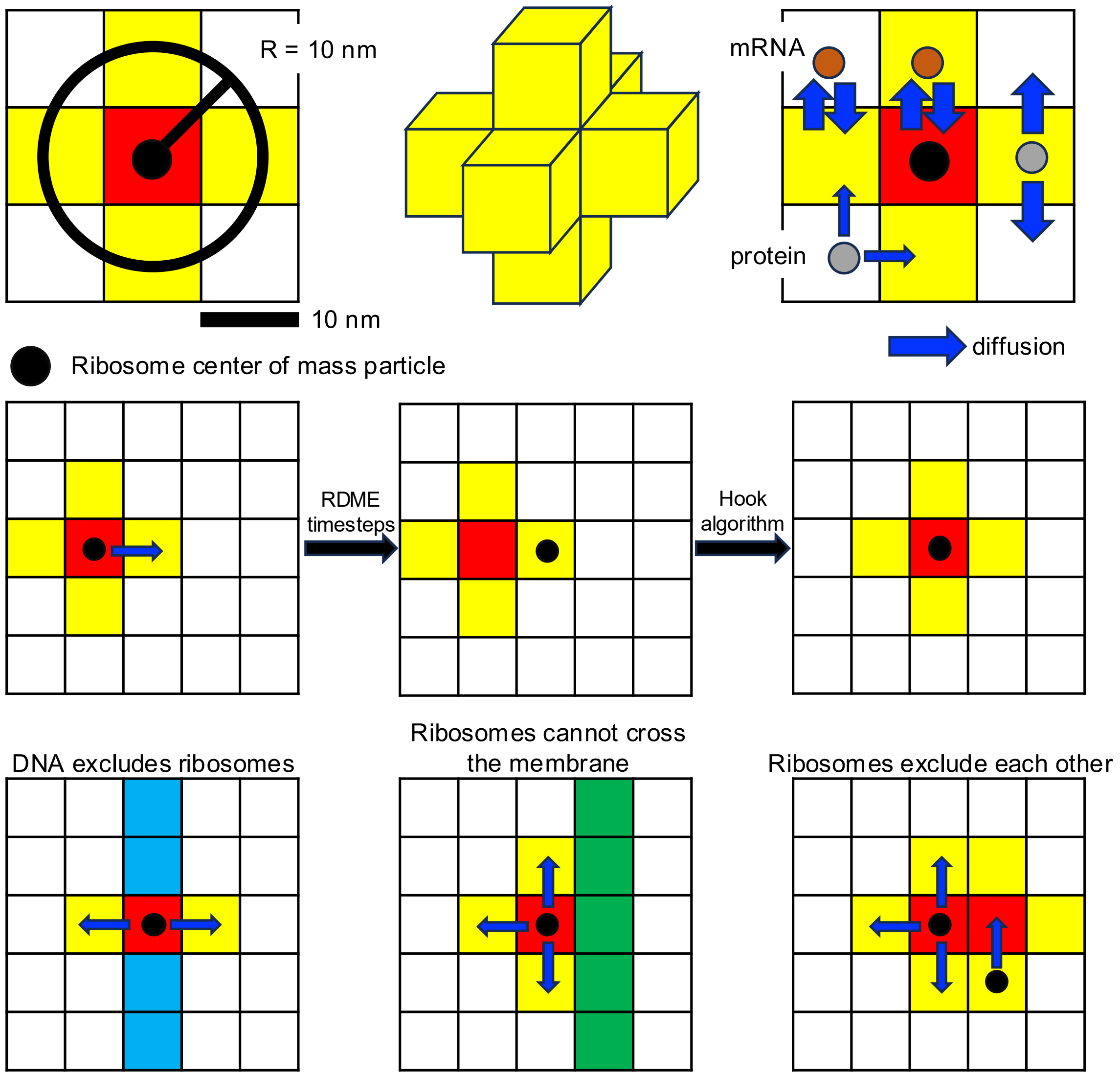


**Figure S1: Projection of the ribosome onto the 10 nm lattice and its diffusion rules.** The ribosome has a radius of 10 nm, resulting in the projections onto the lattice shown into the top left and middle. The top right shows that mRNA can diffuse into and out of ribosomes as their normal cytoplasmic rate so that mRNA can undergo translation initiation. Proteins can diffuse out of ribosomes at their cytoplasmic diffusion rate, but we reduce their diffusion rate into the ribosome because of the excluded volume of the ribosome. The center of mass particle of the ribosome starts at the center lattice site of the ribosome projection and is then allowed to diffuse within the cross. At the communication time, we update the position of the cross to reflect the new position of the center of mass particle.

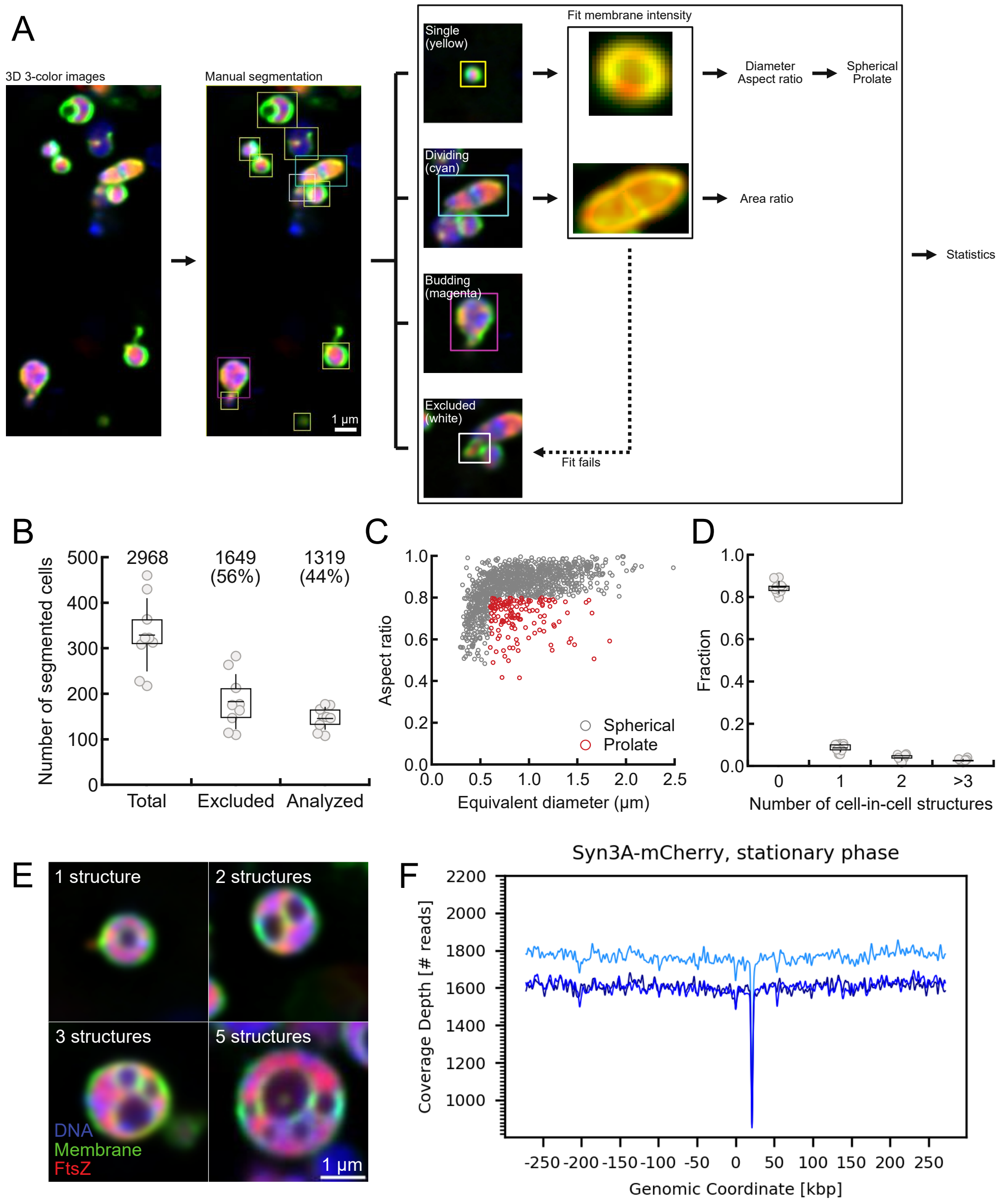


**Figure S2: Imaging statistics and stationary-phase DNA sequencing of JCVI-syn3A/B.** (A) Schematic representation of method to analyze morphologies in fluorescent imaging of JCVI-syn3B-FtsZ:mCherry. Color channels correspond to membrane (green), DNA (blue), and FtsZ (red). (B) Total number of segmented cells, the fraction that were excluded due to cell clusters, and the fraction that were analyzed and included in the morphology statistics. (C) Size and aspect ratio distribution of analyzed single cells. The average cell size was determined to be 0.89±0.37 $\mu$m. The larger size is likely due to sample preparation method and lower spatial resolution in the imaging. Prolate cells were identified by quantifying the aspect ratio of their major and minor axes (see Methods). (D) Less than 20% of cells have "cell-in-cell" structures like the ones shown in (E). These have been observed previously in Syn3A and have been known to form in late exponential phase of growth.[19,32] (F) DNA sequencing was performed in triplicate during the stationary growth phase. The lack of a slope in coverage corresponds to an ori:ter ratio of 1:1, indicating that the cells have stopped replicating. Each dot in a box plot represents a technical replicate (field of imaging) and covers the 25 to 75 percentiles. The means and standard deviations are represented as horizontal and vertical lines, respectively.

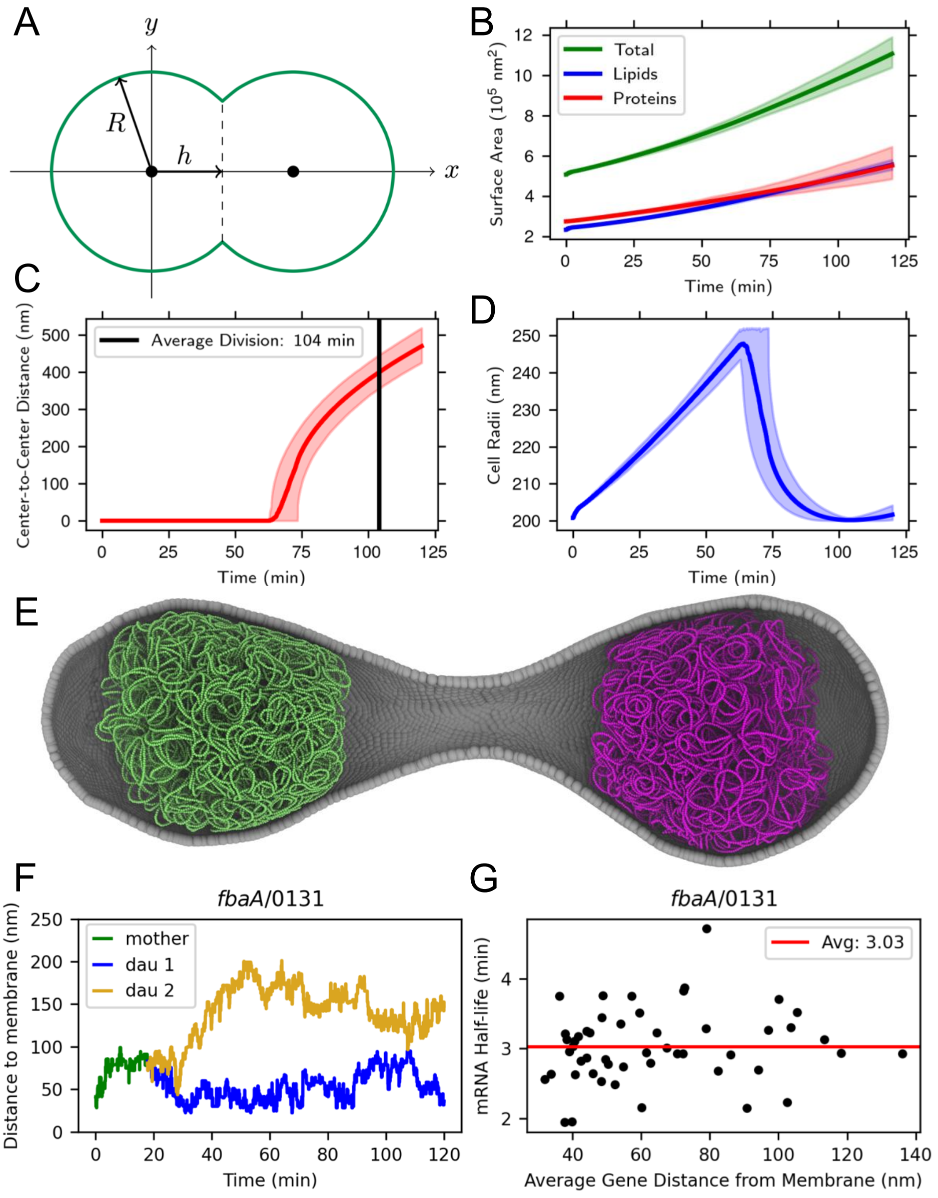


**Figure S3: Spatial components of growth and division.** (A) Schematic of the geometric division model. The dividing cell is treated as two overlapping spheres of radius $R$ whose centers are separated by a center-to-center distance of $2 \times h$. (B) Total surface area and contributions from lipids and membrane proteins. (C) Center-to-center distance ($2 \times h$ from A) calculate from instantaneous cell surface area and volume. (D) Cell radii ($R$ from A) throughout growth and division. The radius increases during spherical growth until division starts once the volume has doubled. In B, C, and D, lines show population average and shaded regions show full range among population. (E) Example of a membrane shape generated using FreeDTS. The long neck was not observed in experimental imaging. Grey particles represent vertices of the triangulated surface. Green and magenta particles represent two chromosomes fitted into the FreeDTS membrane structure. (F) Distance between the transcription start site particle on the RDME lattice for *fbaA* and its nearest membrane lattice site in a single simulated cell. At 20 minutes, the gene is replicated from the mother chromosome to the two daughter chromosomes. (G) Comparison of mRNA half-life to the average distance of the transcription start site particles for *fbaA* to the membrane. Each point represents a single simulated cell.

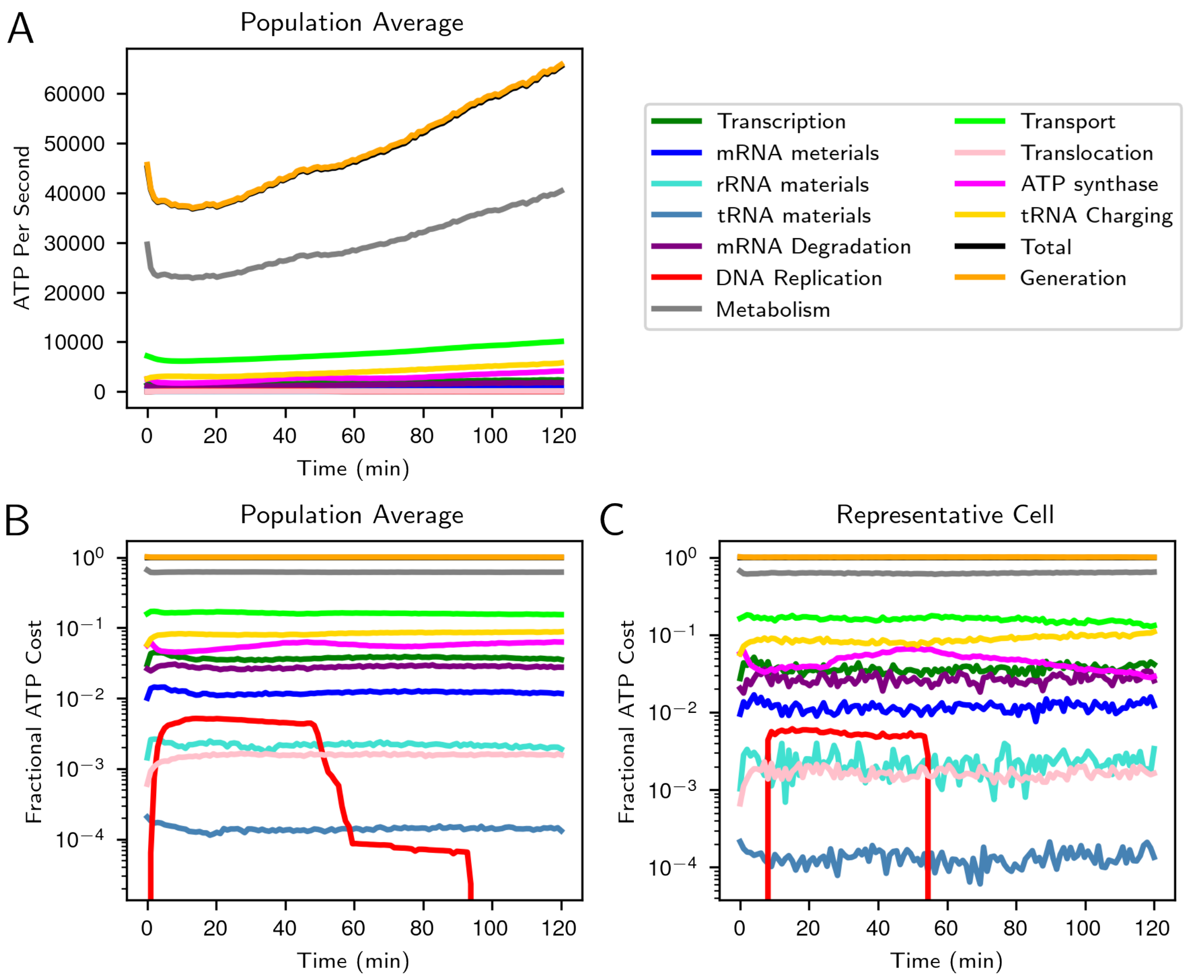


**Figure S4: Cell-wide accounting of all ATP costs.** Costs are averaged among the population (A,B) and for a representative cell (C). The total ATP generated is slightly higher than the total ATP cost to maintain the ATP pool as the cell grows and eventually divides. The cost of DNA replication is only present during replication, and the shoulders observed in the fractional cost plot (B) are a result the variations in the start and end of DNA replication among the population. The long plateau from 60 to 90 minutes is a result of the cell that initiated replication 46 minutes into its cell cycle, resulting in its replication to last until after 90 minutes. While ATP costs are steady on average, there are fluctuations in the costs in a single cell as observed in C.
