## Supplementary material for "Bringing the Genetically Minimal Cell to Life on a Computer in 4D": Key Resources Table

| REAGENT or | SOURCE | IDENTIFIER |
|---|---|---|
| Bacterial and virus strains | | |
| JCVI-syn3A+mCherry | JCVI | N/A |
| JCVI-syn3B+ftsZ:mCherry | JCVI | N/A |
| Chemicals, peptides, and recombinant proteins | | |
| Mycoplasma Broth | BD | Cat#DF0554-17-1 |
| Bacto Tryptone | BD | Cat#211705 |
| Bacto Peptone | BD | Cat#211677 |
| Glucose 20% w/v | Thermo Fisher | Cat#G8270 |
| CMRL 1066 (10X | Thermo Fisher | Cat#21-540-026 |
| Sodium bicarbonate | Thermo Fisher | Cat#S6014 |
| L-glutamine 200 mM | Thermo Fisher | Cat#25030081 |
| Yeast extract | Thermo Fisher | Cat#18180059 |
| TC Yeastolate 2% | Gibco | Cat#255772 |
| Serum (heat | Thermo Fisher | Cat#10828028 |
| Penicillin G | Sigma-Aldrich | Cat#P3032-1MU |
| Phenol red | Sigma-Aldrich | Cat#P0290-100ML |
| KnockOut Serum | ThermoFisher | Cat#10828028 |
| Paraformaldehyde (32%) | Electron Microscopy Sciences | Cat#15680 |
| Phosphate-buffered | Corning | Cat#21-040-CV |
| Poly-D-lysine | Gibco | Cat#A3890401 |
| Hoechst 33342 | Thermo Scientific | Cat#62249 |
| FM1-43FX | Invitrogen | Cat#F35355 |
| Critical commercial assays | | |
| PureLink gDNA | Thermo Fisher | Cat#K182001 |
| Qubit dsDNA HS | Thermo Fisher | Cat#Q33230 |
| Illumina DNA Prep | Illumina | Cat#20060060 |
| Illumina MiSeq | Illumina | Cat#SY-410-1003 |
| Deposited data | | |
| gDNA Sequencing | This study | SRA: SUB15230492 |
| JCVI-syn3A | Breuer et al., 2019 | GenBank: CP016816.2 |
| Mass spectrometry data of Syn3A | Breuer et al., 2019 | MassIVE – Accession Number: 000081687 |
| Proteomics of Syn3A | Breuer et al., 2019 | ProteomeXchange – Accession Number: PXD008159 |
| BRENDA | Chang et al., 2021 | https://www.brenda-enzymes.org/ |
| Software and algorithms | | |
| Lattice Microbes – v2.5 | Luthey-Schulten Lab | https://github.com/Luthey-Schulten-Lab/Lattice_Microbes |
| odecell – v1.0 | Thornburg et al., 2022 | https://github.com/Luthey-Schulten-Lab/odecell |
| FreeDTS version 6.7.2023 | Pezeshkian and Ipsen, 2024 | https://github.com/weria-pezeshkian/FreeDTS |

| | | |
|---|---|---|
| sc_chain_gen version 7.20.2023 | Gilbert et al., 2023 | https://github.com/Luthey-Schulten-Lab/sc_chain_generation |
| LAMMPS version | Thompson et al., | https://www.lammps.org |
| btree_chromo | Gilbert et al., 2023 | https://github.com/Luthey-Schulten-Lab/btree_chromo_gpu |
| Docker version | Docker Inc. | https://www.docker.com/ |
| Apptainer version 1.3.5-1.el8 | Singularity/Apptainer Development Team, Int'l. | https://doi.org/10.5281/zenodo.1310023 |
| VMD – v2.0 alpha | Humphrey et al., 1996 | https://www.ks.uiuc.edu/Research/vmd/ |
| bcl2fastq – v2.20 | Illumina | https://support.illumina.com/sequencing/sequencing_software/bcl2fastq-conversion-software.html |
| Bowtie 2 – v2.5.4 | Langmead and Salzberg, 2012 | https://bowtie-bio.sourceforge.net/bowtie2/index.shtml |
| Samtools – v1.21 | Danecek et al., 2021 | https://github.com/samtools/samtools |
| Other | | |
| 8-well chambered coverslip | Fisher Scientific | Cat#12-565-470 |
